# Efficient Gene Transduction in Pigs and Macaques with the Engineered AAV Vector AAV.GT5 for Hemophilia B Gene Therapy

**DOI:** 10.1101/2022.11.24.517886

**Authors:** Yuji Kashiwakura, Kazuhiro Endo, Atsushi Ugajin, Tomohiro Kikuchi, Shuji Hishikawa, Hitoyasu Nakamura, Yuko Katakai, Nemekhbayar Baatartsogt, Takafumi Hiramoto, Morisada Hayakawa, Nobuhiko Kamoshita, Shoji Yamazaki, Akihiro Kume, Harushi Mori, Naohiro Sata, Yoichi Sakata, Shin-ichi Muramatsu, Tsukasa Ohmori

**Affiliations:** Department of Biochemistry, Jichi Medical University School of Medicine, 3311-1 Yakushiji, Shimotsuke, Tochigi 329-0498, Japan; Department of Surgery, Jichi Medical University, School of Medicine, 3311-1 Yakushiji, Shimotsuke, Tochigi 329-0498, Japan; Center for Development of Advanced Medical Technology, Jichi Medical University, 3311-1 Yakushiji, Shimotsuke, Tochigi 329-0498, Japan; Department of Radiology, Jichi Medical University School of Medicine, 3311-1 Yakushiji, Shimotsuke, Tochigi 329-0498, Japan; The Corporation for Production and Research of Laboratory Primates, 1-16-2 Sakura, Tsukuba, Ibaraki 305-0003, Japan; Center for Gene Therapy Research, Jichi Medical University, 3311-1 Yakushiji, Shimotsuke, Tochigi 329-0498, Japan; Clinical Research Center, Jichi Medical University Hospital, 3311-1 Yakushiji, Shimotsuke, Tochigi 329-0498, Japan; Devision of Neurological Gene Therapy, Center for Open Innovation, Jichi Medical University, 3311-1 Yakushiji, Shimotsuke, Tochigi 329-0498, Japan; Center for Gene and Cell Therapy, The Institute of Medical Science, The University of Tokyo, Tokyo 108-8639, Japan

## Abstract

Gene therapy for hemophilia using adeno-associated virus (AAV) vectors allows long-term coagulation factor expression. We examined the potential of a novel engineered liver-tropic AAV3B-based vector AAV.GT5 for hemophilia B gene therapy. *In vitro* transduction with AAV.GT5 in human hepatocytes was more than 100 times higher than with AAV-Spark100, while *in vivo* transduction efficacy into the liver and the increase in coagulation factor IX (FIX) antigen following intravenous injection of these vectors were similar in PXB mice (chimeric mice with a humanized liver) and macaques. The discrepancy was due to the low recovery and short half-life of AAV.GT5 in blood, depending on the positive charge of the heparin-binding site in the original AAV3B. The intra-hepatic vascular administration of AAV.GT5, but not AAV-Spark100, enhanced vector transduction into the liver and reduced vector distribution to the kidney in pigs. In macaques, the intra-hepatic artery injection of AAV.GT5 yielded a comparable increase in FIX antigen with a one-third dosage of peripheral venous administration. Two of four macaques who received AAV.GT5 intravenously did not develop neutralizing antibodies (NAbs) against AAV.GT5, while AAV-Spark100 induced serotype-specific NAbs in all four macaques. The NAb produced after the administration was relatively specific to the serotype and less responsive to the other serotype. As a result, the administration of AAV.GT5 successfully boosted FIX expression in one animal previously given AAV-Spark100. Thus, AAV.GT5 has different biodistribution and immunogenic characteristics compared with AAV-Spark100, and the intra-hepatic vascular administration may lessen the vector dose and avoid vector distribution to other organs.

**Key Points:**

- The AAV.GT5 vector has a strong transduction efficacy in human hepatocytes but has a faster clearance after systemic administration.
- Intra-hepatic vascular administration of the AAV.GT5 vector is an effective liver transduction method for hemophilia gene therapy.

## Introduction

Gene therapy with AAV vectors is attracting attention as a promising treatment for hemophilia, and several clinical trials on gene therapy are currently ongoing.^1^ A single vector administration increases coagulation activity for the long-term, drastically reducing the need for coagulation factor concentrates.^2–7^ AAV vectors possess only a DNA cassette for functional coagulation factor expression, replacing viral Rep and Cap genes between inverted terminal repeat (ITR) sequences. After systemic administration, AAV vectors are internalized into the target cells via the cognate receptors and subsequently form episomal double-stranded concatemers, where the exogenous gene can be expressed over a long period.^8^

Various serotypes of AAV have been used in gene therapy for hemophilia, including naturally occurring AAVs such as AAV8 and AAV5,^2,4^ and engineered AAVs. AAV-Spark100, a modified vector closely related to rh10 in sequence, has shown therapeutic efficacy in hemophilia B clinical trials at doses as low as 5 × 10^11^ vg/kg.^3^ A study showing the therapeutic efficacy of modified BBM-H901, which is similar to AAV1 and AAV6, are also reported.^9^ AAV-LK03 was developed as an engineered serotype with high gene transfer to human hepatocytes^10^ and used for clinical trials of hemophilia A.^7^ Very recently, results of human clinical trials of AAVS3, a hybrid AAV vector of AAV8 and AAV3B, showed high efficacy with low vector doses.^5^

Because the efficiency of gene transfer to the liver hepatocytes varies with serotype, vector doses vary considerably among hemophilia clinical trials, ranging from 2 × 10^11^ to 6 × 10^13^ vg/kg.^1^ It has been reported that a high-dose AAV vector can cause several adverse events due to immune responses. Thus, reducing the vector dose is one important aspect of ensuring the safety of AAV-based gene therapy.^11–13^ Another problem with AAV vectors is the patient’s immune status against AAV. Like the naturally occurring anti-AAV-neutralizing antibodies (NAbs) attenuates AAV-mediated gene therapy,^14,15^ induction of NAbs after an AAV vector administration deprives a chance to re-administrate the same serotype of the AAV vector. Thus, strategies must be developed for efficient gene transfer to the liver at lower doses and re-administration of AAV vectors evading NAbs.

We have recently developed a novel engineered AAV vector AAV.GT5, based on AAV3B sequence.^16^ AAV.GT5 has a high transduction efficiency in human hepatocytes and is relatively insensitive to NAbs found in blood donors.^16^ Therefore, AAV.GT5 may become a vector for efficient hemophilia gene therapy. We investigated the characteristics and efficacy of AAV.GT5 and another engineered AAV, AAV-Spark100, to determine whether AAV.GT5 could provide an effective gene therapy for hemophilia.

## Methods

### Cell culture

The detailed information is described in Supplemental methods.

### Vector construction

Codon-optimized human coagulation factor IX (hFIX) Padua (R338L) minigene (insertion of truncated *F9* intron between exon 1 and exon2) with reduced CpG sequences was synthesized by Thermo Fisher Scientific (Waltham, MA). The DNA fragment consisting of HCRhAAT (an enhancer element of the hepatic control region of the Apo E/C1 gene and the human antitrypsin) promoter, hFIX-Padua minigene, and SV40 polyA was inserted between ITRs derived from AAV2. The capsid sequence of pRC5 (Takara Bio, Shiga, Japan) was replaced by AAV.GT5 and AAV-Spark100 for production of each AAV serotype.

### AAV vector production

The methods for AAV production are described in Supplemental methods. The quality of the AAV vector was examined by sedimentation velocity analytical ultracentrifugation at U-Medico Inc. (Osaka, Japan). We confirmed that AAV vectors produced by our purification method contained more than 83.5% full particles (Supplemental figure S1).

### AAV vector transduction and measurement of firefly luciferase *in vitro*

The detailed methods are described in Supplemental methods.

### Animal experimentation

All experimental animal procedures were approved by The Institutional Animal Care and Concern Committee of Jichi Medical University (permission number: 19029-07, 20023-01, 20054-02, 20051-06), Shin Nippon Biomedical Laboratories (SBL712-003), and Tsukuba Primate Research Center (DSR03-15). Animal care was conducted following the committee’s guidelines and ARRIVE guidelines.^17,18^ The detailed methods for the animal experiments are described in Supplemental methods.

FIX-deficient mice (*B6.129P2-F9^tm1Dws^*) were obtained from The Jackson Laboratory (Sacramento, CA, USA). PXB mice, chimeric mice with a humanized liver repopulated by human hepatocytes, were obtained from Phenix Bio (Hiroshima, Japan). The PXB mice had 83%–96% engraftment of human hepatocytes used in the experiments. CB17/IcrJcl-*Prkdc^scid^* (SCID mouse) were purchased from CLEA Japan (Tokyo, Japan).

The intravenous injection of AAV vector into *Macaca fascicularis* (macaque) was performed at Shin Nippon Biomedical Laboratories (Kagoshima, Japan). The characteristics of macaques are shown in Supplemental table S1. AAV vector expressing hFIX-Padua was intravenously administrated via saphenous vein for 5 minutes. One macaque administered with AAV.GT5 was considered not to receive the AAV vector efficiently because we could not detect the predicted AAV genomes in serum 60 minutes after administration (#2). Hence, we excluded the data of the macaque (#2) from the analysis. The intra-hepatic artery AAV vector administration into macaques was performed at Tsukuba Primate Research Center, National Institutes of Biomedical Innovation, Health and Nutrition (Ibaraki, Japan). We confirmed the insertion of the catheter into the proper hepatic artery by contrast imaging and then administrated the AAV vector for 5 minutes. We intramuscularly administrated 1 mg/kg of prednisolone daily for 56 days, and tapered thereafter [0.5 mg/kg/day (57–63 days), 0.3 mg/kg/day (64–70 days), 0.2 mg/kg/day (71–77 days), 0.1 mg/kg/day (78–84 days)] to reduce the vector immunogenicity.

Porcine experiments were conducted at the Center for Development of Advanced Medical Technology (CDAMTec) at Jichi Medical University. Microminipigs were obtained from Fuji Micra (Shizuoka, Japan). The characteristics of the animals are shown in Supplemental table S2. Systemic administration of AAV vectors was performed via ear vein. The intra-hepatic artery AAV vector administration was performed, as in the case of macaques. Intraportal administration of AAV was done, as described previously.^19^ The same dose (1 mg/kg/day) of prophylactic prednisolone was administered intravenously only for 33 days.

### Anti-AAV NAb assay

We measured anti-AAV NAb titer in sera from 216 patients with hemophilia and 100 healthy volunteers. The Institutional Review Board at Jichi Medical University approved the study protocols (permission number: A19-108), and we obtained written informed consent from all the participants. The study was registered in UMIN-CTR (UMIN-CTR: UMIN000039069).^20^ The detailed methods are described in Supplemental methods.

### Measurement of FIX activity and antigen

hFIX activity (FIX:C) and hFIX antigen (FIX:Ag) were measured using chromogenic assay or sandwich ELISA, as described in Supplemental methods.

### Isolation of anti-hFIX antibodies from macaque plasma with inhibitors

We used a specific anti-hFIX antibody from a macaque that developed a FIX inhibitor (#7) to detect hFIX:Ag in macaque plasma. The detailed method is described in Supplemental methods.

### Quantitative polymerase chain reaction (qPCR) and amplicon sequencing

AAV genome and mRNA expression levels in organs were measured by quantitative polymerase chain reaction (qPCR) and reverse transcription-directed qPCR (RT-qPCR), respectively. When indicated, PCR amplicons were subjected to 300 pair-end read sequencing using Illumina MiSeq (Illumina, San Diego, CA). The detailed methods are described in Supplemental methods.

## Results

### Efficient transgene expression in human hepatocytes by AAV.GT5 *in vitro*

We compared the *in vitro* transduction efficiency of AAV.GT5 with that of parental AAV3B vector. Luciferase expression by AAV.GT5 was significantly higher than that by AAV3B in human hepatocellular carcinoma cell line Huh-7 (Supplemental figure S2). We compared *in vitro* transduction efficiency among AAV5, AAV-Spark100, and AAV.GT5 in human or mouse hepatocytes. AAV-Spark100 is the engineered AAV serotype that has succeeded in treating hemophilia B at a low dose,^3^ and AAV5 is a natural AAV serotype used in some hemophilia B gene therapy trials.^4^ AAV.GT5 showed robust transgene expression in human primary hepatocytes as well as Huh-7 and HepG2 (Supplemental figure S2), as reported previously.^16^ Transduction efficiency of human primary hepatocytes with AAV.GT5 was more than 100 times higher than AAV-Spark100 and AAV5 (Supplemental figure S2). Conversely, AAV-Spark100 showed the most efficient transduction in murine hepatocyte cell line TLR3 (Supplemental figure S2).

### The transgene expression in PXB mice after intravenous injection of AAV.GT5 and AAV-Spark100

We compared the *in vivo* liver transduction efficiency of AAV.GT5 with that of AAV-Spark100. We constructed AAV vectors expressing luciferase gene driven by CAG promoter and then intravenously injected them into PXB mice (animals with humanized liver).^21^ Both vectors showed comparable levels of luciferase expression in the liver assessed by *in vivo* imaging and AAV genomes in the liver (Figure 1). *In vivo* experiments showed efficient liver transduction with AAV-Spark100 in SCID mice (Supplemental figure S3). We constructed an AAV vector harboring hFIX with Padua mutation (R338L) driven by liver-specific HCRhAAT promoter (HCRhAAT-hFIX-Padua). Intravenous injection of the AAV.GT5 in FIX-deficient mice showed an increase in plasma FIX:Ag and FIX:C in a dose-dependent manner (Supplemental figure S4). The ratio of FIX:C to FIX:Ag in FIX Padua was about 8–10-fold (Supplemental figure S4), as reported previously.^22^ We further compared the increase in plasma FIX levels by intravenous injection of AAV.GT5 and AAV-Spark100 harboring HCRhAAT-hFIX-Padua in PXB mice. We administered the vector to PXB mice and assessed the plasma level of FIX:C. A similar level of increase in FIX:C in PXB mice was obtained by AAV.GT5 and AAV-Spark100 (Figure 1D). The AAV genome and hFIX mRNA in the liver did not differ significantly between the groups (Figures 1E and F).

**Figure 1.**
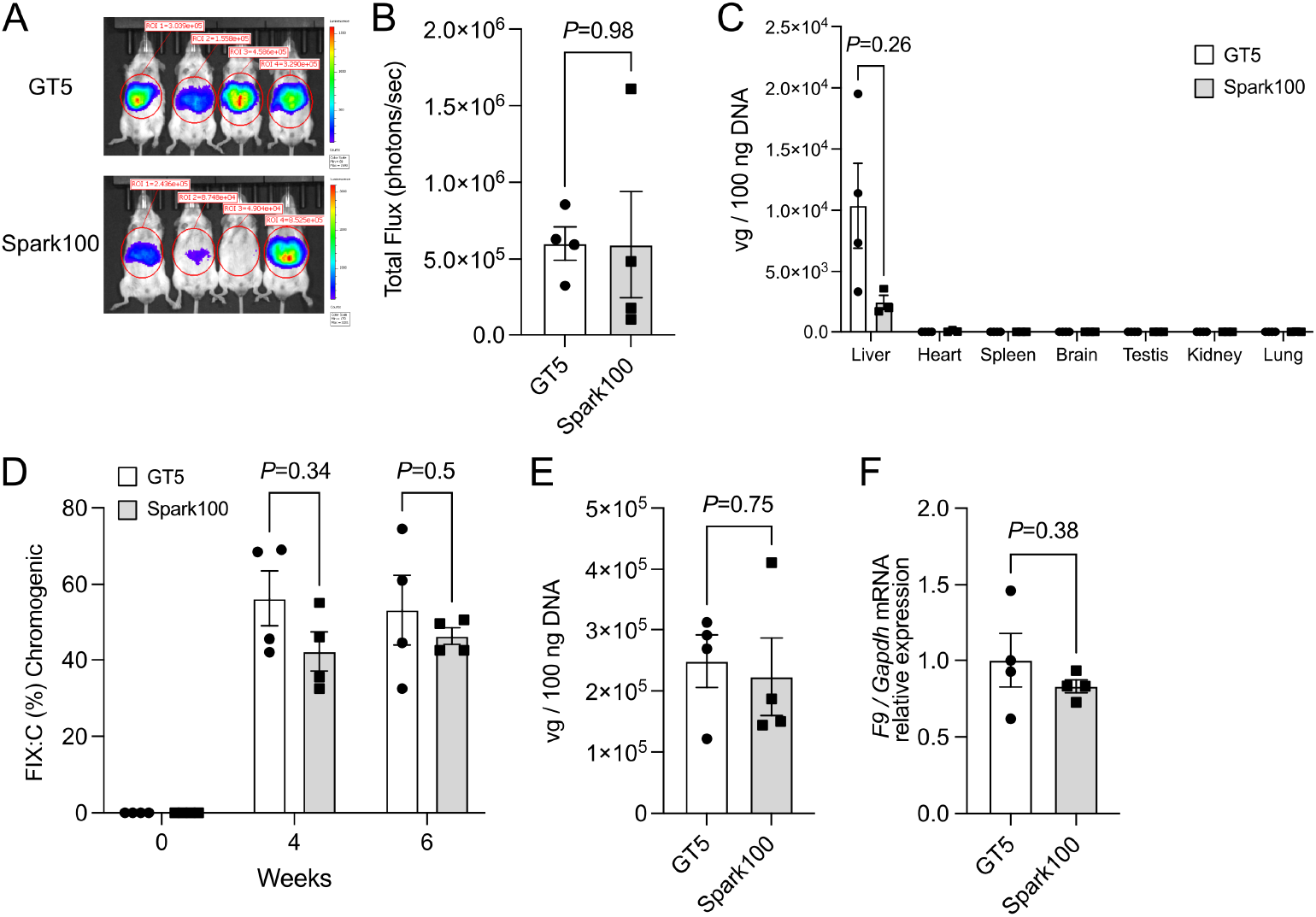
The transgene expression in PXB mice by intravenous injection of AAV.GT5 and AAV-Spark100. (A, B, C) AAV.GT5 or AAV-Spark100 harboring luciferase gene driven by CAG promoter were injected intravenously into PXB mice (1 × 10^12^ vg/kg). Photons transmitted through the body were analyzed using an IVIS Imaging System. (A) Representative data of IVIS imaging system at 4 weeks after the vector administration. (B) Quantitative data of each mouse were expressed as photon units (photons/seconds). Values are presented as mean ± SEM (n = 4). (C) AAV genomes in organs were determined by quantitative PCR 8 weeks after the vector administration. Values are presented as mean ± SEM (n = 4 in AAV.GT5 and n = 3 in AAV-Spark100). (D, E) AAV.GT5 or AAV-Spark100 harboring human coagulation factor IX with Padua mutation (hFIX-Padua) gene driven by HCRhAAT promoter were injected intravenously into PXB mice (2 × 10^12^ vg/kg). (D) The plasma was obtained at 0, 4, and 6 weeks after the vector injection. The increase in hFIX activity (FIX:C) from the baseline (0 weeks) was observed. Values are presented as mean ± SEM (n = 4). (E) AAV genome in the liver was determined by quantitative PCR 7 weeks after the vector administration. Values are presented as mean ± SEM (n = 4). (F) hFIX mRNA expression in the liver was assessed using real-time RT-PCR and was expressed as the fold increase in the *F9/GAPDH* ratio. Values are presented as mean ± SEM (n = 4). Statistical analysis between the two groups was performed by Student *t-*test. Actual *P* values are described in the figure.

### The transgene expression in nonhuman primates by systemic intravenous injection of AAV.GT5 and AAV-Spark100

We assessed the transgene expression in macaques following intravenous injection of AAV.GT5 and AAV-Spark100. We selected the macaques without NAbs against AAV.GT5 or AAV-Spark100 in respective vector administration (Supplemental table S1). We administered AAV-Spark100 or AAV.GT5 harboring HCRhAAT-FIX Padua at 1 × 10^12^ vg/kg intravenously into macaques. A significant increase in FIX:Ag and FIX:C was observed after the vector administration. The plasma levels of FIX:Ag and FIX:C reached the maximum at 7–14 days after administration and gradually decreased over time (Figure 2). The levels of FIX:Ag and increase in FIX:C did not significantly differ between AAV.GT5 and AAV-Spark100 (Figure 2). After the vector administration, the anti-hFIX neutralizing antibody (coagulation factor inhibitor) was developed in 2 macaques (#4 and #7, Figure 3). The serum chemistry test results, including liver enzymes, are shown in Supplemental figure S5. The biodistribution study of AAV showed efficient transduction in the liver by both vectors. The levels of the AAV genome in the liver were not significantly different between the groups (Figure 2). Trace amounts of the AAV genome were detected in other organs, but there was no significant difference between AAV.GT5 and AAV-Spark100 (Supplemental figure S6).

**Figure 2.**
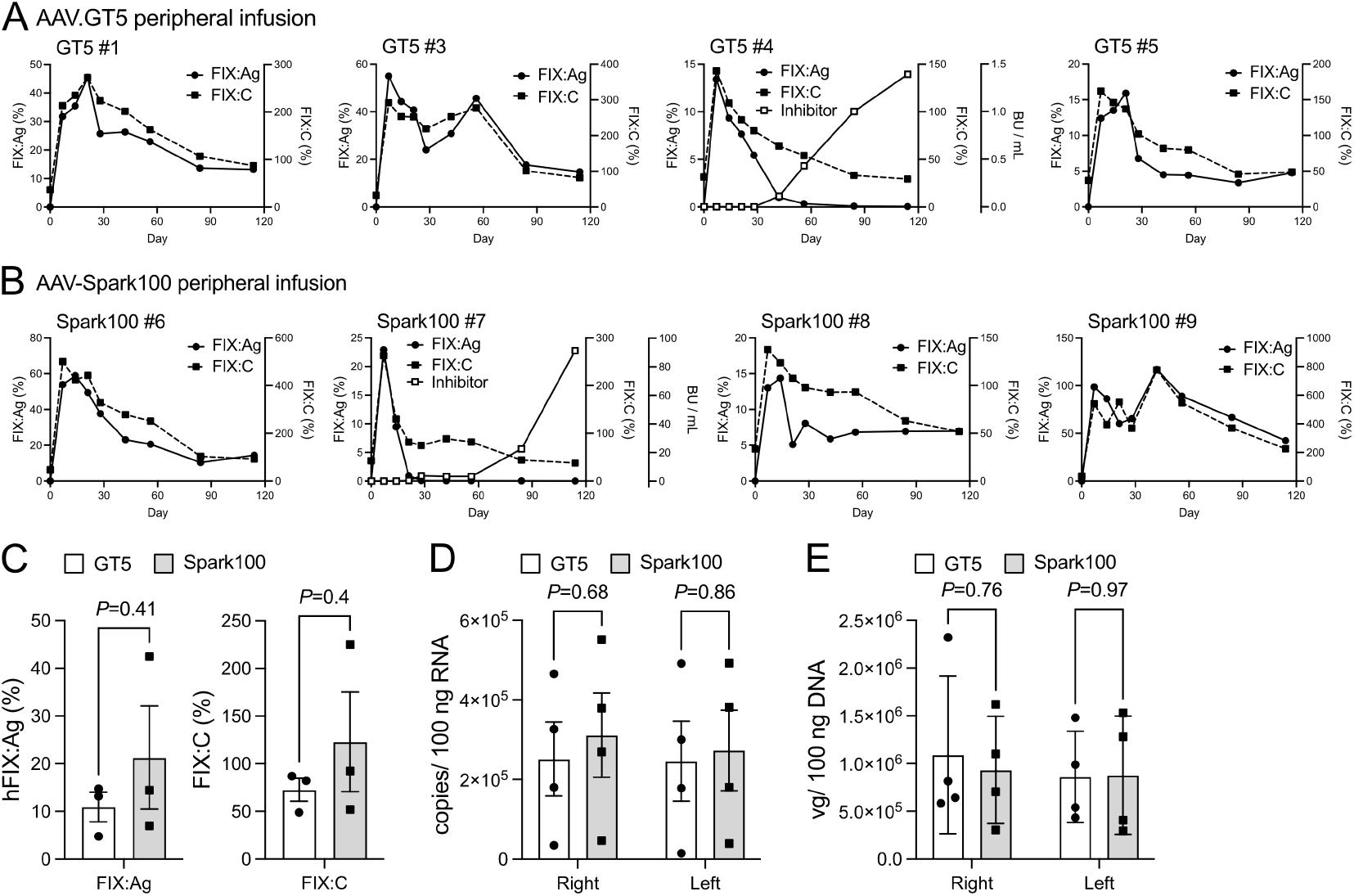
The plasma levels of hFIX and AAV genome in macaques after intravenous injection of AAV.GT5 and AAV-Spark100. (A, B) AAV.GT5 (A) or AAV-Spark100 (B) harboring human coagulation factor IX with Padua mutation (hFIX-Padua) gene driven by HCRhAAT promoter were injected intravenously into cynomolgus macaques (1 × 10^12^ vg/kg). The changes in plasma levels of hFIX antigen (FIX:Ag) and increase in hFIX activity (FIX:C) in each macaque were observed. FIX neutralizing antibody was developed in two macaques (AAV.GT5 in #4 and AAV-Spark100 in #7). (C) Comparison of plasma levels of FIX:Ag 114 days after the vector administration between AAV.GT5 and AAV-Spark100. Values are presented as mean ± SEM (three macaques without hFIX inhibitors in each group). (D) hFIX mRNA expression in the liver was assessed using real-time RT-PCR and was expressed as an *F9* copy number in 100 ng RNA. Values are presented as mean ± SEM (n = 4). (E) The AAV genome in livers was determined by quantitative PCR 114 days after the vector administration. Values are presented as mean ± SEM (n = 4). Statistical analysis between the two groups was performed by Student *t-*test. Actual *P* values are described in the figure.

**Figure 3.**
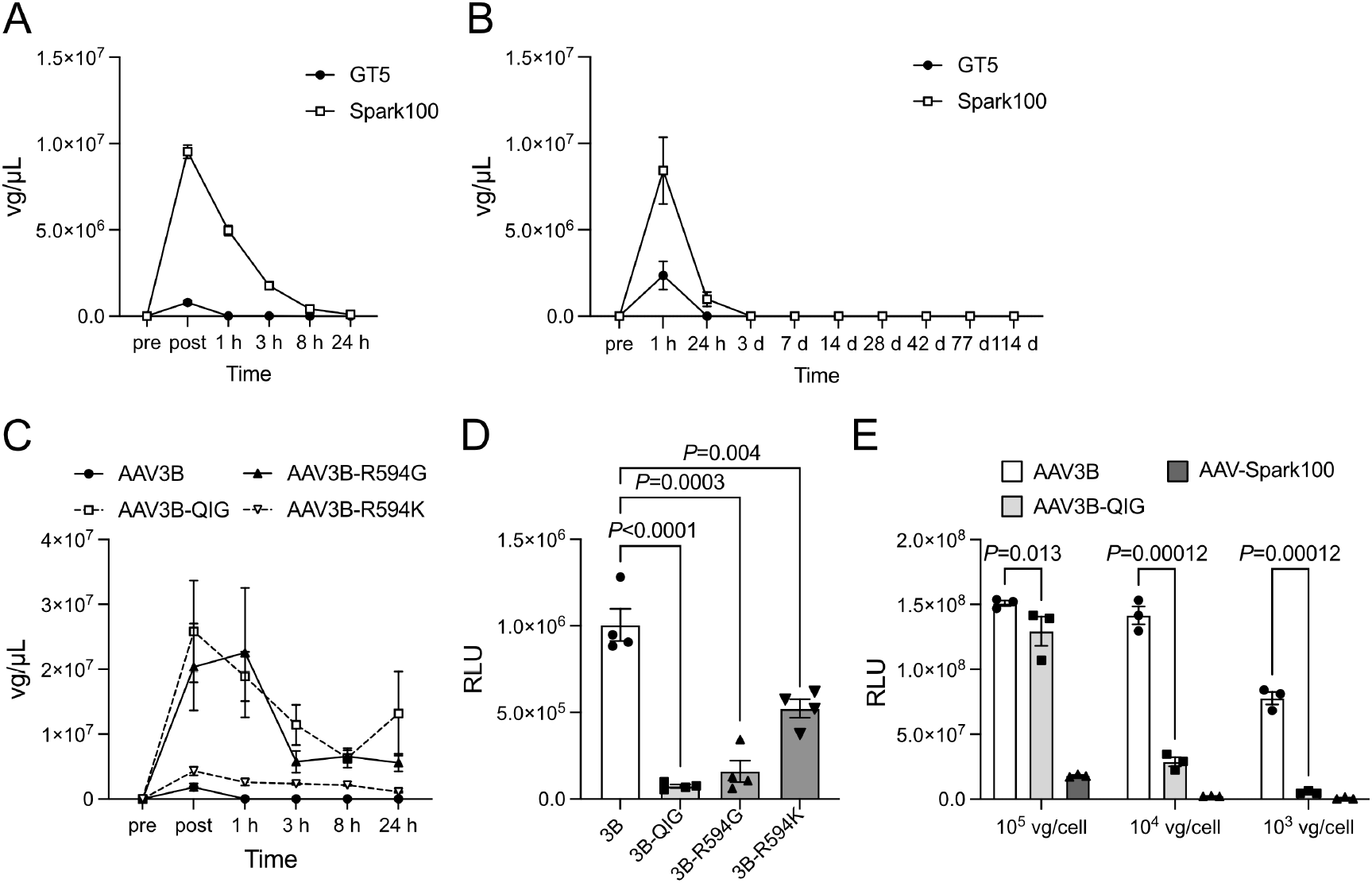
Differences in vector clearance between AAV.GT5 and AAV-Spark100. (A) AAV.GT5 or AAV-Spark100 harboring luciferase gene driven by CAG promoter were injected intravenously into C57BL/6 mice (1 × 10^12^ vg/kg). Values are presented as mean ± SEM (n = 4). The changes of plasma AAV genomes were determined by quantitative PCR and expressed as AAV vector genome in plasma. (B) AAV.GT5 or AAV-Spark100 harboring human coagulation factor IX with Padua mutation (hFIX-Padua) gene driven by HCRhAAT promoter were injected intravenously into macaques (1 × 10^12^ vg/kg). Values are presented as mean ± SEM (n = 4). The changes in plasma AAV genomes were determined by quantitative PCR and expressed as AAV vector genomes in plasma. (C, D) AAV3B vector without or with mutation(s) at heparin-binding site [QIG (T592Q, T593I, and R594G), R594G, or R594K] harboring secreting type of NanoLuc were injected intravenously into C57BL/6 mice (1 × 10^12^ vg/kg). (C) The changes of plasma AAV genomes were determined by quantitative PCR and expressed as AAV vector genome in plasma. Values are presented as mean ± SEM (n = 4). (D) Plasma levels of NanoLuc activity were determined by luminometer 4 weeks after the vector administration. Values are presented as mean ± SEM (n = 4). (E) Transduction of Huh-7 cells with AAV3B vector without (AAV3B) or with mutation(s) in heparin-binding site [QIG (T592Q, T593I, and R594G)] or AAV-Spark100 harboring secreting type of NanoLuc. Transductions with AAV vectors were assessed by NanoLuc activity in the supernatant. Values are presented as mean ± SEM (n = 3). Statistical analysis between the two groups was analyzed by Student *t-*test. Actual *P* values are described in the figure.

### Difference in vector kinetics between AAV.GT5 and AAV-Spark100

Although AAV.GT5 exhibited strong transduction efficiency in hepatocytes *in vitro;* the *in vivo* transduction efficacy did not outrun compared with AAV-Spark100 following intravenous injection. We hypothesized that the discrepancy between *in vitro* and *in vivo* transduction efficacies is due to the faster disappearance of AAV.GT5 from the circulation after systemic delivery. Hence, we compared the kinetics of the vector genome in mouse blood after intravenous injection. The recovery and blood retention of AAV.GT5 was significantly poor than that of AAV-Spark100 (Figure 3A). The recovery rate soon after the intravenous injection of AAV.GT5 was 5.0%–10.6%, while that of AAV-Spark100 was 75.3%–90.2%. Serum concentrations of AAV genome in macaques treated with AAV.GT5 was also significantly lower than in macaques treated with AAV-Spark100 (Figure 3B). A kinetics study showed that serum AAV.GT5 quickly disappeared, unlike AAV-Spark100 (Figure 3B). Comparing the amino acid sequence of the capsid proteins suggested that the difference may attribute to the heparin-binding site of the parental AAV3B capsid. When the heparin-binding site of the AAV3B amino acid sequence (TTR) was changed into the AAV8 sequence (QIG), the recovery and half-life of the vector were significantly improved (Figure 3C). The replacement of the amino acid “R (arginine)” in the heparin-binding site by “G (glycine),” but not by “K (Lysine),” also clearly prolonged the half-time (Figure 3C), suggesting the positive charge in the heparin-binding site is determinant for the short half-life of AAV3B-based vectors. These replacements of the heparin-binding site reduced the vector transduction *in vitro* and *in vivo* (Figure 3D and E).

### Efficient transduction in the liver with intra-hepatic vascular administration of AAV.GT5 in pigs and nonhuman primates

We considered the low recovery and fast clearance of AAV.GT5 *in vivo* may be due to the binding of AAV.GT5 to negatively charged substances, including heparin sulfate through the heparin-binding site, and offset the advantage of efficient transduction of liver hepatocytes in *in vivo* gene therapy, particularly when given into peripheral veins. Administration to large vessels directly perfusing the liver, such as the portal vein and hepatic artery, may take better advantage of the high transduction efficiency of AAV.GT5 into hepatocytes. We employed microminipigs to compare the transduction efficacies of the AAV vector through different injection routes because of easy access to hepatic vessels. We confirmed the efficient transduction with AAV.GT5 in primary porcine hepatocytes *in vitro* (Supplemental figure S7). We compared the transduction efficiency of AAV vectors through a peripheral vein with those through the portal vein. We approached the left portal vein through the supra mesenteric vein with the catheter (Video 1). The plasma levels of FIX:Ag and FIX:C in pigs treated with AAV-Spark100 harboring HCRhAAT-FIX Padua were not different after intravenous injection and portal vein injection (Figure 4A).

**Figure 4.**
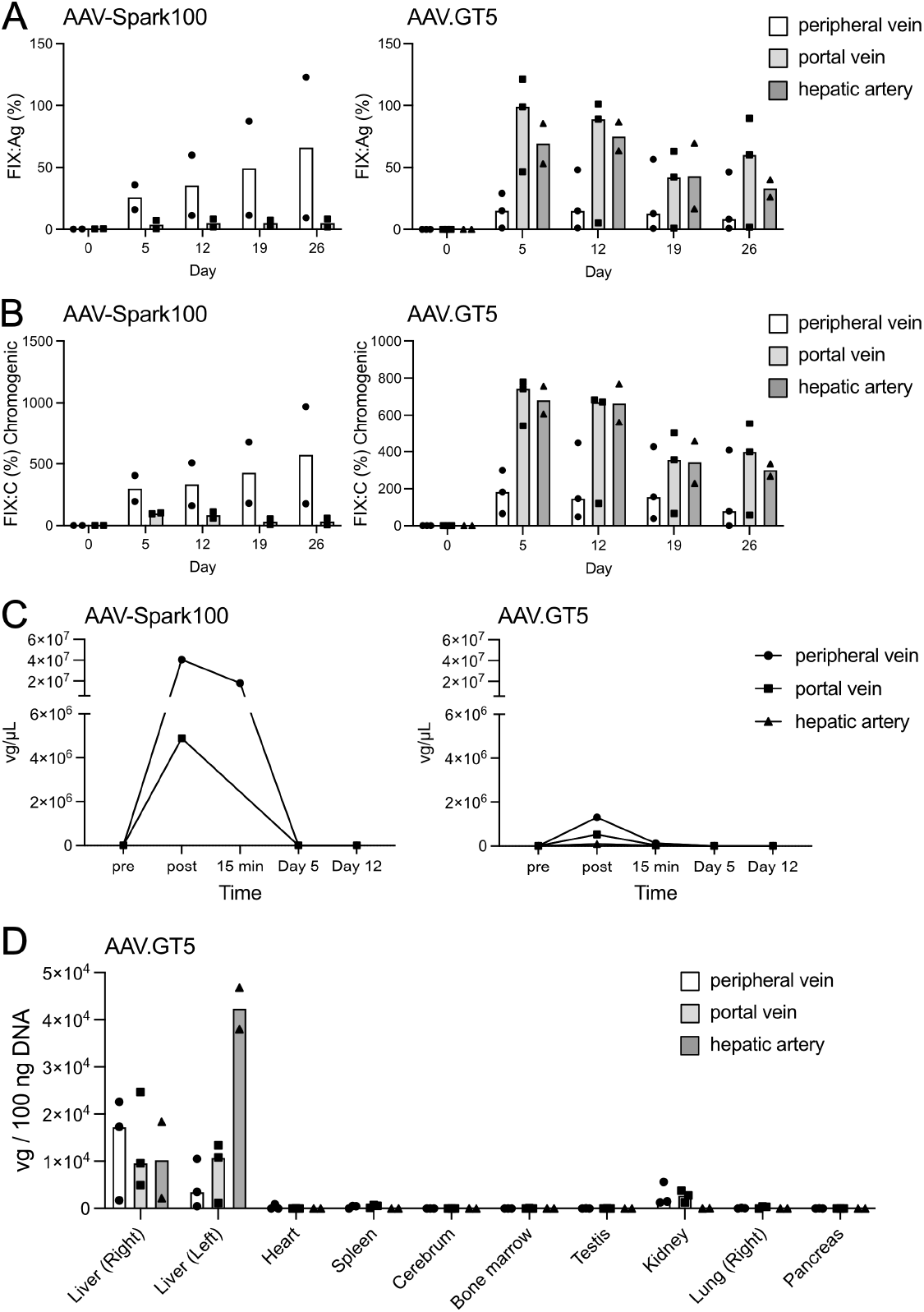
The difference in transduction efficacy with AAV.GT5 among the administration routes in pigs. AAV.GT5 or AAV-Spark100 harboring human coagulation factor IX with Padua mutation (hFIX-Padua) gene driven by HCRhAAT promoter were administrated into male microminipig through a peripheral vein, left portal vein, or hepatic artery (1 × 10^12^ vg/kg). (A) The changes in plasma FIX:Ag after the administration in the pig treated with AAV-Spark100 or AAV.GT5 through the peripheral vein, left portal vein, or hepatic artery. Bar, median (AAV-Spark100: n = 2, AAV.GT5: n = 3 in a peripheral vein and portal vein; n = 2 in a hepatic artery). (B) The changes in plasma FIX:C after the administration in the pigs treated with AAV-Spark100 or AAV.GT5 through the peripheral vein, portal vein, or hepatic artery. Bar, median (AAV-Spark100: n = 2, AAV.GT5: n = 3 in a peripheral vein and portal vein; n = 2 in a hepatic artery). (C) The changes in plasma AAV genome in pigs treated with AAV-Spark100 or AAV.GT5. Values are presented as median (AAV-Spark100: n = 2, AAV.GT5: n = 3 in a peripheral vein and portal vein; n = 2 in a hepatic artery). (D) AAV genome in organs obtained from the pigs treated with AAV.GT5 was determined by quantitative PCR 33 days after the vector administration. Bar, median (n = 3 in peripheral and portal veins; n = 2 in a hepatic artery).

The intraportal injection of AAV.GT5 showed a significant increase in plasma FIX levels than that after intravenous injection (Figure 4A and B, and Supplemental figure S8). We assessed the effect of intra-hepatic artery injection of AAV.GT5 in pigs as it is a less invasive procedure. We inserted the catheter through the femoral artery and directly assessed the proper hepatic artery through the celiac artery (Video 2). We confirmed similar high transduction efficiencies of AAV.GT5 through intra-hepatic artery injection (Figure 4A and B, and Supplemental figure S8). We confirmed the higher recovery and retention of AAV-Spark100 in blood compared with that of AAV.GT5 (Figure 4C). Interestingly, administration via a hepatic artery or portal vein resulted in poor recovery in pigs (Figure 4C). After liver-directed vector administration, we did not observe any abnormality in blood chemistry, including liver enzymes (Supplemental Figure S9). We obtained a high AAV genome in the left liver lobe after intra-hepatic artery injection, whereas the distribution of the AAV genome in the kidney was low (Figure 4D).

Based on the above experiments, we supposed the intra-hepatic artery injection of AAV.GT5 enhanced liver transduction and was thus able to reduce the vector dose. We next administrated AAV.GT5 harboring HCRhAAT-FIX Padua through the hepatic artery at one-third vector dose (3 × 10^11^ vg/kg) of the peripheral vein administration in macaques (Video 3). As expected, it significantly increased plasma FIX:Ag and FIX:C (Figure 5). The levels of FIX:Ag, FIX:C, and AAV genome in the liver were similar after peripheral intravenous administration at 3 times the dose (1 × 10^12^ vg/kg) (Figure 5). The course of laboratory test parameters after intra-hepatic artery injection of the vector is shown in Supplemental figure S10.

**Figure 5.**
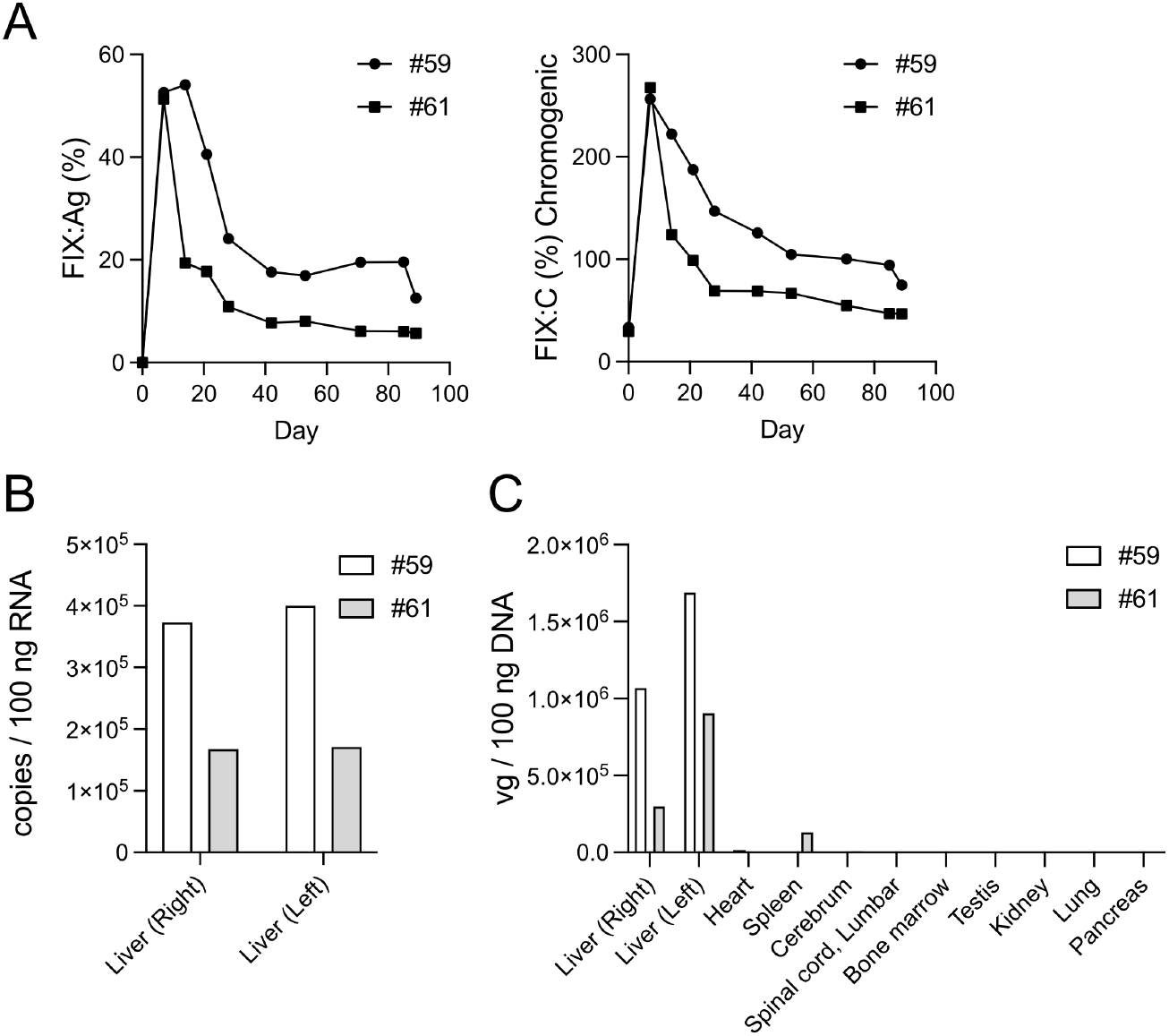
Efficient transduction of the liver with AAV.GT5 through hepatic artery injection in macaques. AAV.GT5 harboring human coagulation factor IX with Padua mutation (hFIX-Padua) gene driven by HCRhAAT promoter were injected into two male macaques through the hepatic artery (3 × 10^11^ vg/kg). (A) The changes in plasma levels of hFIX antigen (FIX:Ag) and increase in hFIX activity (FIX:C) in each macaque were observed. (B) hFIX mRNA expression in the liver was assessed using real-time RT-PCR and was expressed as *F9* gene copies in 100 ng RNA. (C) AAV genomes in organs treated were determined by quantitative PCR 89 days after the vector administration.

### Threshold of NAb titer to inhibit vector transduction

The anti-AAV NAbs abolish the treatment benefit of systemic AAV injection, but little is known about the NAb level that affects the efficacy. We examined the threshold of NAb to inhibit vector transduction by systemic injection with AAV.GT5 or AAV-Spark100 *in vivo.* We injected different concentrations of human immunoglobulin in C57BL/6 mice and then administrated AAV.GT5 or AAV-Spark100 expressing secNanoLuc at three doses (Supplemental figure S11). A low titer of NAb was enough to inhibit the low AAV vector transduction dose significantly. However, a higher dose could evade the inhibitory effect of the low dose of NAb (Supplemental figure S11). The threshold of NAbs to inhibit vector transduction at the same vector dose (5 × 10^12^ vg/kg) was much higher in AAV.GT5, compared with AAV-Spark100 (AAV.GT5, IC50: 7.15; AAV-Spark100, IC50: 3.75) (Supplemental figure S11).

### Comparison of NAb titer in healthy volunteers and patients with hemophilia in Japan

We measured the seroprevalence of anti-AAV NAbs against AAV.GT5 and AAV-Spark100 in 100 healthy volunteers and 216 patients with hemophilia in Japan. The prevalence and the titer of anti-AAV NAbs against AAV.GT5 and AAV-Spark100 were not significantly different (Figure 6A and B). NAb titers against AAVGT5 and AAV-Spark100 showed a strong cross-correlation (Supplemental figure S12). There was no significant difference in the seroprevalence of NAbs between AAV3B and AAV.GT5, but NAb titers were significantly lower in AAV.GT5 in the subjects with NAb titers less than 1:1000 (Supplemental figure S13).

**Figure 6.**
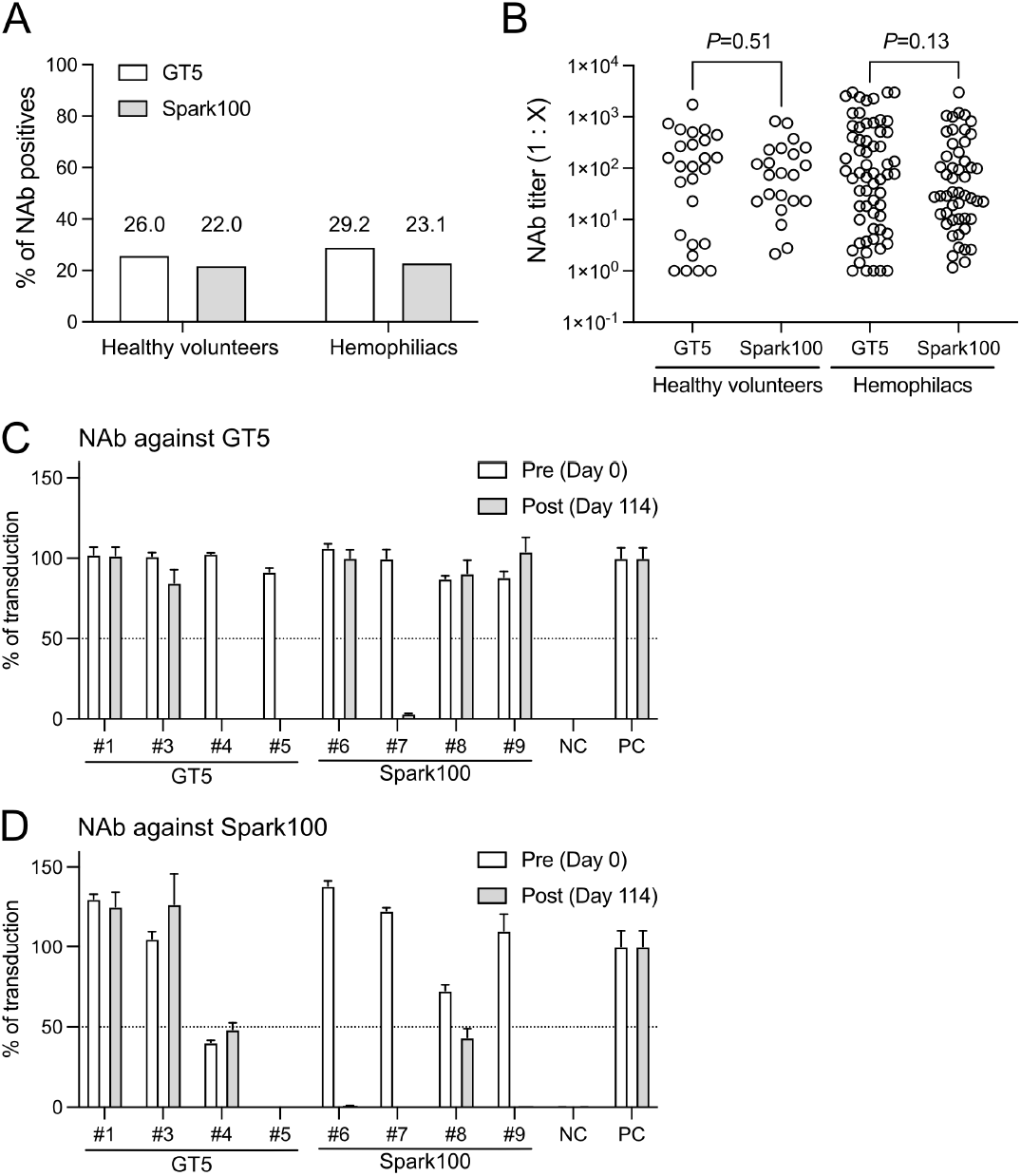
Neutralizing antibody against AAV.GT5 and AAV-Spark100. (A) The seroprevalence of neutralizing antibodies (NAbs) against AAV.GT5 and AAV-Spark100 in healthy volunteers (n = 100) and patients with hemophilia (n = 216). The statistical difference among serotypes was analyzed using the *C* test. (B) The NAb titer against AAV.GT5 and AAV-Spark100 in healthy volunteers and patients with hemophilia. Statistical differences among serotypes were analyzed using the Mann-Whitney *U* test. Actual *P* values were described in the Figure. (C, D) AAV.GT5 or AAV-Spark100 harboring human coagulation factor IX with Padua mutation (hFIX-Padua) gene driven by HCRhAAT promoter were injected intravenously into macaques (1 × 10^12^ vg/kg). The inhibition of transduction efficacy with AAV.GT5 (C) and AAV-Spark100 (D) by the addition of serum (1:1) obtained at pre-administration (day 0) or post-administration (day 114). The data are presented as mean ± SD of triplicated samples. A NAb positive was considered when infection was inhibited by 50% or more.

### Emergence of NAb in nonhuman primates after vector administration

We analyzed the emergence of NAb against AAV after AAV vector administration in nonhuman primates. We measured NAb against AAV.GT5 and AAV-Spark100 before and 114 days after the vector injection. The treatment with AAV-Spark100 induced the Nabs against AAV-Spark100 in all macaques (Figure 6D). Only one macaque treated with AAV-Spark100 developed Nabs against AAV.GT5 (#7) (Figure 6C). Two of four macaques treated with AAV.GT5 did not seroconvert after the vector administration (Figure 6C). NAbs against AAV-Spark100 were not changed in the macaques treated with AAV.GT5 (Figure 6D).

### Booster effect of AAV.GT5 in pig previously treated with AAV-Spark100

We examined the efficacy of the second challenge of the AAV vector in pigs. Of the two animals previously treated with intravenous administration of AAV-Spark100 harboring HCRhAAT-FIX Padua (Figure 4), one did not develop the NAbs against AAV.GT5, the other developed Nab against AAV.GT5 (1:15.8). To explore the possibility of the second dosing, we administered AAV.GT5 harboring HCRhAAT-FIX into the animal without NAbs through the hepatic artery. We employed wild-type FIX, not Padua mutation, to discriminate the transduction efficacy of 1^st^ and 2^nd^ administration by the next-generation sequencer. The re-administration with AAV.GT5 significantly increased the plasma FIX:Ag (Figure 7A). The next-generation sequencing of the liver DNA and mRNA expression of hFIX showed the efficient transduction of the liver by re-administration of AAV.GT5 through the hepatic artery (Figure 7B and C).

**Figure 7.**
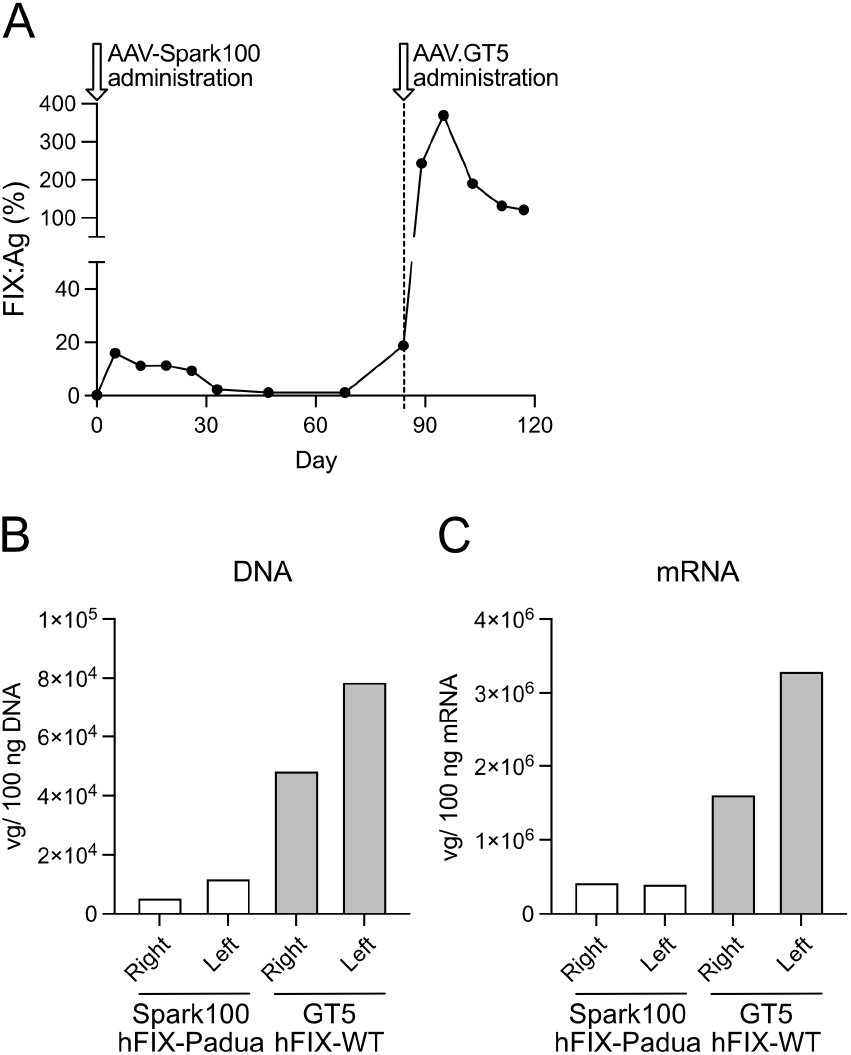
Re-administration of AAV.GT5 in a pig treated with AAV-Spark100. A microminipig was first intravenously treated with AAV-Spark100 harboring human coagulation factor IX with Padua mutation (hFIX-Padua) gene driven by HCRhAAT promoter (1 × 10^12^ vg/kg) (#21-006 in Supplemental figure S8). At 84 days after the first injection, AAV.GT5 harboring wild type hFIX (hFIX-WT) gene driven by HCRhAAT promoter were administrated through the hepatic artery. (A) The changes in plasma levels of hFIX antigen (FIX:Ag) were shown. (B, C) mRNA expression (B) AAV genome (C) in the liver was determined by quantitative PCR 117 days after the first administration. The expressions of the transgene (mRNA) and transduction efficacy (AAV genome) with first and second administration were distinguished by next-generation sequencing.

## Discussion

It is important to ensure the safety of gene therapy to decrease the immune response and off-targeting to other organs by reducing the vector dose in AAV vector-mediated gene therapy. Minimizing the immunogenicity of the vector is beneficial for vector re-administration. We have developed the new AAV3B-based engineered AAV serotype AAV.GT5, with a substitution of three amino acids in the antigenic epitope of AAV.^19^ In this study, we compared the gene transfer efficiency of AAV.GT5 into the liver with that of AAV-Spark100 to prove the efficacy in hemophilia B gene therapy. AAV.GT5 has strong transduction efficiency in human hepatocytes while it is rapidly eliminated in blood after intravenous injection. Taking advantage of these properties, the modification of the administration route has been shown to result in high transduction efficacy in the liver at low vector doses *in vivo.* AAV.GT5 may be less immunogenic with regard to NAb induction, which usually occurs after the AAV vector administration.

The intra-hepatic vascular administration of AAV.GT5 may reduce vector dose and restrict the organ distribution of vectors in liver-directed gene therapy. We found that the positive charge of the heparin-binding site derived from AAV3B is important for this property. The heparin-binding site, the most strong positively charged region on the AAV3B surface that is not conserved in other serotypes,^23^ affected vector clearance, possibly through the binding to negatively charged substances such as heparin sulfate on endothelial cells. On the other hand, the enhancement of vector transduction by the intra-hepatic vascular administration was not observed in the case with AAV-Spark100, which has a high-recovery rate and long half-life due to the lack of the heparin-binding site. Although AAV vectors have been considered safe with low immunogenicity, recent reports have indicated adverse reactions such as liver injury, thrombotic microangiopathy, and degeneration of posterior ganglia in animal models due to the induction of immune responses when administered at high doses.^11,24–28^ Considering the systemic adverse reactions due to vectors, reducing the dose of vectors administered in the systemic injection is desirable. Arterial administration may prevent the distribution of vectors to organs other than the liver, because hepatic arterial administration results in the rapid disappearance of AAV in the blood. In addition, the production costs may lessen when the amount of vector decreases.

We found the possibility of low immunogenicity of AAV.GT5 after the vector injection. Systemic administration of AAV produces NAbs against the administered AAV serotype without exception. However, we could not observe a serotype-specific NAb production in two of four macaques intravenously treated with AAV.GT5. This study is the first to report no NAb development after vector administration in large animal models, although the macaques had received immunosuppression with prednisolone. Under the same conditions, all macaques treated with AAV-Spark100 developed NAbs. The two macaques with developed antibodies after AAV.GT5 administration had possessed NAbs against AAV-Spark100 before the administration. NAb production against AAV.GT5 may unlikely occur if the macaque is naïve to infection against any AAV. We recently assessed the prevalence of NAbs against nine AAV serotypes in 100 healthy subjects and 216 patients with hemophilia in Japan.^20^ We found that 69% of Japanese patients with hemophilia were negative for Nabs against all AAVs, which is considered a naïve situation for AAV infection. It is very vital to confirm the NAb production after AAV.GT5 administration in clinical trials in the future.

One of the important challenges of AAV-mediated gene therapy is re-administration. NAbs against the administrated AAV are produced after vector injection strongly abolished the second administration of the same AAV vector. AAV.GT5 might be repeatedly injected if the NAbs would not occur after the first challenge. The status of NAbs to the other AAV serotype was virtually unchanged, suggesting the NAbs after the vector injection was relatively specific to the serotypes. Recent data from long-term observation of dogs with hemophilia have also shown the emergence of specific NAbs based on the administered AAV serotype.^29^ This is in sharp contrast to the cross-reactivity of NAbs induced by natural infection in patients with hemophilia, which inhibited multiple AAV serotypes.^20^ We have shown that AAV.GT5 vector can transduce a transgene to the liver in pigs treated with AAV-Spark100. It is important to have several serotypes of gene therapy products for a single disease, assuming that the efficacy of the vector is not achieved or gradually disappears.

This study has several limitations. First, we observed a decrease in FIX levels from 7~14 days after administration, despite using a codon-optimized FIX sequence with reduced CpG. The apparent transaminase elevation was not observed, but it is probably due to the elimination of AAV-infected liver cells by an immune response. Although prednisolone was used as immunosuppression in large animal studies, it may be necessary to use immunosuppressive agents in actual clinical practice more effectively. We obtained higher peak FIX antigen levels in macaque and porcine experiments than we expected before the experiments. The reduction of vector dose may reduce the number of AAV entering one transduced hepatocyte, resulting in a reduced immune response and sustained therapeutic effects. Secondly, the number of experiments using large animals is limited in this study. In some cases, the number of experiments was only performed on two animals; thus, statistical differences were not obtained. It is necessary to increase the number of cases before human clinical trials. We reported that the heparin-binding site is associated with the recovery and clearance of the vector. However, the detailed mechanisms and where the vector is consumed *in vivo* are unknown. Finally, we only showed the results of animal experiments, even though we used animal species that are close to humans. We must prove the therapeutic efficacy of AAV.GT5 for liver transduction in a human clinical trial.

In conclusion, we reported the therapeutic potential of a new AAV3B-based engineered vector AAV.GT5 for hemophilia gene therapy. We have demonstrated that the administration of GT5 via the hepatic artery could reduce the dose of the vector and systemic dissemination of the vectors by taking advantage of the properties of AAV.GT5. This may lead to further improvement in the safety and efficacy of gene therapy using AAV vectors. AAV.GT5 can be applied in clinical trials for gene therapy for various genetic diseases, including hemophilia, in the future.

## Acknowledgments

This work was supported by AMED [JP18pc0101030]. Optima XE-90 was subsidized by JKA through its promotion funds from KEIRIN RACE. We thank Yaeko Suto, Mika Kishimoto, Tamaki Aoki, Sachiyo Kamimura, Mai Hayashi, Yuiko Ogihara, Nagako Sekiya, Noguchi Tomoko, Hiromi Ozaki, Hiroko Hayakawa, Mika Ito, and Naomi Takino of Jichi Medical University for their technical assistance. We would like to acknowledge Dr. Tomoyuki Abe and Dr. Yoshimitsu Izawa of Jichi Medical University for their assistance in porcine surgery and autopsy. We are also grateful to Dr. Azusa Nagao (Ogikubo Hospital), Dr. Kagehiro Amano (Tokyo Medical University), Dr. Nobuaki Suzuki (Nagoya University), Dr. Tadashi Matsushita (Nagoya University), Dr. Akihiro Sawada (Hyogo College of Medicine), Dr. Satoshi Higasa (Hyogo College of Medicine), Dr. Naoya Yamazaki (Hiroshima University), Dr. Teruhisa Fujii (Hiroshima University), Dr. Taemi Ogura (Shizuoka Children’s Hospital), Dr. Hideyuki Takedani (IMSUT Hospital, The University of Tokyo), Dr. Masashi Taki (St. Marianna University), Dr. Takeshi Matsumoto (Mie University), Dr. Jun Yamanouchi (Ehime University), Dr. Michio Sakai (Munakata Suikokai General Hospital), Dr. Masako Nishikawa (The University of Tokyo), Dr. Yutaka Yatomi (The University of Tokyo), Dr. Koji Yada (Nara Medical University), and Dr. Keiji Nogami (Nara Medical University), for recruiting patients for the NAb assay.

## Disclosures

All authors declare no competing financial interests.

## Author contributions

Y.K. designed the study, performed the experiments, analyzed the data, and wrote the manuscript. K.E., A.U., T.K., S.H., H.N., Y.K., N.B. T.H., M.H., and N.K., S.Y., H.M., S.M. designed the study, performed the experiments, and revised the manuscript. A.K., N.S., and Y.S. designed the study and revised the manuscript. T.O. designed the study, performed the experiments, analyzed the data, and wrote the manuscript. All authors approved the final version of the manuscript.

## Notes

### Competing Interest Statement

The authors have declared no competing interest.

