## Supplement for "Efficient Gene Transduction in Pigs and Macaques with the Engineered AAV Vector AAV.GT5 for Hemophilia B Gene Therapy"

**Short title:** The Engineered Vector AAV.GT5 for Hemophilia B

**Supplemental methods**
**Supplemental tables S1–S3**
**Supplemental figures S1–S13**
**Videos 1–3**

### Supplemental methods

#### Cell culture

AAVpro 293T cells (Takara Bio, Shiga, Japan) and Huh-7 cells (Riken BRC, Ibaraki, Japan) were cultured in Dulbecco modified Eagle Medium (DMEM, Sigma Aldrich, Saint Louis, MO) supplemented with 10% fetal bovine serum (FBS, Thermo Fisher Scientific, Waltham, MA) and 2 mM L-glutamine (Thermo Fisher Scientific). The murine immortalized hepatocytes TLR3 cells (JRCB Cell Bank, Osaka, Japan) were cultured in DMEM supplemented with 2% FBS, 10 ng/mL of human epidermal growth factor (Thermo Fisher Scientific), and ITS supplement (Thermo Fisher Scientific). Cryopreserved human primary hepatocytes and minipig primary hepatocytes were purchased from Kaly-Cell (Plobsheim, France) and Sekisui XenoTech (Kansas City, KS), respectively.

#### AAV vector transduction and measurement of firefly luciferase *in vitro*

The cell lines were seeded at a density of $1 \times 10^4$ cells in each well of 96-well plates coated with collagen type I (Cellmartrix Type I-A, Nitta Gelatin, Osaka, Japan) on the day before the transduction. The primary hepatocytes were seeded at a density of $5 \times 10^4$ cells in each well of 96-well plates coated with collagen type I. A vial of vector stock was thawed and diluted with DMEM containing 5% FBS immediately before the transduction experiment and was directly added to each well. The cells were lysed with 50 µL of 1× Passive Lysis Buffer (Promega, Madison, WI) 48 hours after the transduction and immediately stored in a deep freezer. Following a previously described study by Baatartsogt et al.[1] the luciferase activity was measured by a luminometer (Centro LB 960, BERTHOLD Technologies, Bad Wildbad, Germany).

#### AAV vector production

The AAV genes were packaged by triple plasmid transfection of AAVpro 293T cells to generate the AAV vector (helper-free system), as described previously.[2] AAV vectors were purified from the transfected cells 72 hours after the transduction using the ultracentrifugation method, as described previously.[3] Titration of recombinant AAV vectors was performed by quantitative PCR targeting the SV40 polyA sequence.[1] The quality of the AAV vector was examined by sedimentation velocity analytical ultracentrifugation at U-Medico Inc. (Osaka, Japan). We confirmed that AAV vectors produced by our purification method contained more than 83.5% full particles (Supplemental figure S1).

#### Animal experimentation

All experimental animal procedures were approved by The Institutional Animal Care and Concern Committee of Jichi Medical University (permission number: 19029-07, 20023-01, 20054-02, 20051-06), Shin Nippon Biomedical Laboratories (SBL712-003), and Tsukuba

Primate Research Center (DSR03-15). Animal care was conducted following the committee's guidelines and ARRIVE guidelines.[4,5]

Coagulation factor IX (FIX)-deficient mice (B6.129P2-*F9*$^{tm1Dws}$) were obtained from The Jackson Laboratory (Sacramento, CA, USA). PXB-mice, chimeric mice with a humanized liver repopulated by human hepatocytes, were obtained from Phenix Bio (Hiroshima, Japan). The PXB-mice had 83%–96% engraftment of human hepatocytes used in the experiments. CB17/IcrJcl-*Prkdc*$^{scid}$ (SCID mouse) were purchased from CLEA Japan (Tokyo, Japan). Mice were maintained in isolators in the specific pathogen-free facility of Jichi Medical University at 23°C ± 3°C with 12:12 hours light/dark cycle. Mice were anesthetized with isoflurane (1%–3%) to obtain plasma samples, and the blood sample was drawn from the jugular vein with a 29G micro-syringe (TERUMO, Tokyo, Japan) containing 1/10 (volume/volume) sodium citrate (Harasawa Pharmaceutical, Tokyo, Japan). Platelet-poor plasma was isolated by centrifugation and then frozen and stored at −80°C until the analysis. AAV vector (100 μL) was administrated intravenously through the jugular vein. The mice were anesthetized with isoflurane to analyze the luciferase expression in the liver *in vivo* and then received the luciferin substrate (3 mg/body weight) intraperitoneally. Photons transmitted through the body were analyzed using an IVIS® Imaging System and Living Image software (Xenogen Corp., Alameda, CA). Quantitative data were expressed as photon units (photons/second).

The intravenous injection of the AAV vector into *Macaca fascicularis* was performed at Shin Nippon Biomedical Laboratories (SNBL, Kagoshima, Japan). We selected nine male macaques without anti-AAV Nabs against AAV-Spark100 or AAV.GT5. The characteristics of monkeys are shown in Supplemental table S1. AAV vector expressing human FIX Padua was intravenously administrated via a saphenous vein for 5 minutes. One monkey treated with AAV.GT5 could not efficiently administrate the AAV vector because of failure to detect the AAV genome in serum 60 minutes after the administration (#2). Hence, we excluded the data of the monkey (#2) from the analysis. The intra-hepatic artery AAV vector administration into *Macaca fascicularis* was performed at Tsukuba Primate Research Center, National Institutes of Biomedical Innovation, Health and Nutrition (Ibaraki, Japan). We confirmed the insertion of the catheter into the proper hepatic artery by contrast imaging and then administrated the AAV vector for 5 minutes. To reduce the vector immunogenicity, prednisolone was intramuscularly administered at 1 mg/kg/day for 56 days, and then gradually reduced [0.5 mg/kg/day (57–63 days), 0.3 mg/kg/day (64–70 days), 0.2 mg/kg/day (71–77 days), 0.1 mg/kg/day (78–84 days)].

Pig experiments were conducted at the Center for Development of Advanced Medical Technology (CDAMTec) at Jichi Medical University. Microminipigs were obtained from Fuji Micra (Shizuoka, Japan). The characteristics of pigs are shown in Supplemental table S2. Intravenous systemic administration of AAV vectors was performed via ear vein. The intra-hepatic artery AAV vector administration was performed, as in the case of *Macaca* *fascicularis*. Intraportal administration of AAV was done, as described previously.[6] Prednisolone was intravenously administered at 1 mg/kg/day for 33 days.

**Anti-AAV NAb assay**

We measured anti-AAV Nab titer against AAV-Spark100 and AAV.GT5 in sera from 216 patients with hemophilia and 100 healthy volunteers. The Institutional Review Board at Jichi Medical University approved the study protocols (permission number: A19-108), and we obtained written informed consent from all the participants. The study was registered in UMIN-CTR (UMIN-CTR: UMIN000039069).[7] The measurement of Nab was essentially performed as described previously.[8] Briefly, the sera were heat-inactivated (56°C, 30 min), diluted with FBS to 1:1–1:3,000, and then incubated with an AAV vector expressing luciferase under the control of CAG promoter for 1 hour at 37°C. The mixture was added in triplicates to a 96-well plate seeded with Huh-7 cells at a multiplicity of infection (MOI) of 1500. After 48 hours of incubation, cells were lysed with 50 µL of 1× Passive Lysis Buffer (Promega) and stored in a deep freezer. The luciferase activity in the lysates was measured as described previously.[1] We estimated antibody titers that neutralized 50% vector transduction ($ND_{50}$) by nonlinear regression using GraphPad Prism 9 (GraphPad, San Diego, CA). The correlation of NAb titers between AAV.GT5 and AAV-Spark100 were examined by Spearman's rank correlation coefficient using GraphPad Prism 9 (GraphPad). We created the topological phylogenetic tree of AAV serotypes using the maximum likelihood method with Molecular Evolutionary Genetics Analysis 11 software.[9]

**Measurement of FIX activity and antigen**

Human FIX (hFIX) (FIX:C) was measured by Revohem FIX Chromogenic, a blood coagulation FIX measurement kit with an automated coagulation analyzer (Sysmex CS-1600 analyzer; Sysmex, Kobe, Japan). hFIX antigen (FIX:Ag) in mouse and porcine plasma was measured by sandwich ELISA kit (VisuLizet™ Factor IX Antigen Kit, Affinity Biologicals, Ancaster, ON, Canada), according to the manufacturer's recommendation. The isolated antibody against hFIX was coated in 96-well plates (1 µg/mL) overnight at 4°C. After blocking with 5% casein in PBS for 1 hour at room temperature, samples were incubated in PBS containing 1% casein and 0.1% Triton X-100 for 1 hour at 37°C. After washing with PBS containing 0.1% Triton X-100, the bound FIX antigen was detected with the anti-hFIX antibody conjugated with horseradish peroxidase (Affinity Biologicals) using ABTS microwell peroxidase substrate (Seracare, Milford, MA).

**Isolation of anti-hFIX antibodies from monkey plasma with inhibitors**

We developed a polyclonal monkey anti-hFIX antibody from the serum of one monkey with FIX inhibitor (#7) to detect hFIX:Ag in monkey plasma. The serum obtained from the monkey inhibited human FIX:C, but not monkey FIX:C (data not shown), suggesting the emergence of hFIX-specific antibodies. Immunoglobulin G (IgG) was isolated from the monkey serum using Melon$^{TM}$ Gel IgG Purification Kit (Thermo Fisher Scientific). The antibody specifically bound to hFIX was isolated using column chromatography (AminoLink Plus Immobilization Kit, Thermo Fisher Scientific) conjugated with recombinant hFIX (Benefix, Pfizer, New York, NY).

**Quantitative polymerase chain reaction (qPCR) and amplicon sequencing**

Genomic DNA and RNA were extracted by DNeasy Blood & Tissue Kit (QIAGEN, Venlo, Netherlands) and RNeasy Mini kit (QIAGEN), respectively. The RNA samples were reverse-transcribed using a PrimeScript RT Reagent kit (Takara Bio). Quantitative polymerase chain reaction (qPCR) was performed using THUNDERBIRD Probe qPCR Mix (TOYOBO, Osaka, Japan) or THUNDERBIRD™ SYBR qPCR Mix (TOYOBO) on QuantStudio 12K Flex (Thermo Fisher Scientific). The quantification of the AAV genome into genomic DNA was measured by the quantification of the codon-optimized hFIX Padua sequence. The AAV genome in organs obtained from PXB mice was measured by quantifying the luciferase gene. mRNA expression levels of ectopic human codon-optimized FIX was expressed as copy number in 100 ng RNA. When indicated, amplicon sequencing was performed at Bioengineering Lab (Kanagawa, Japan). PCR amplicons were subjected to 300 pair-end read sequencing using Illumina MiSeq (Illumina, San Diego, CA). The frequencies of DNA sequences were analyzed using CRISPResso2 (https://crispresso.pinellolab.partners.org). The primer and probe sequences are described in Supplemental table S3.

**Supplemental table S1. Cynomolgus monkeys included in this study**

| Number | Sex | Individual number | Age | Body weight (kg) | Vector used | Vector dose (vg/kg) | Administration route | NAb titer against GT5 | NAb titer against Spark100 | MAX of hFIX Ag (%) | Note |
|---|---|---|---|---|---|---|---|---|---|---|---|
| #1 | male | SC0405045 | 15 | 7.75 | AAV.GT5 | $1 \times 10^{12}$ | peripheral vein | neg | neg | 45.54 | |
| #2 | male | SC1301139 | 7 | 7.82 | AAV.GT5 | $1 \times 10^{12}$ | peripheral vein | neg | 10 | 9.46 | Failed to administer |
| #3 | male | SC1304309 | 7 | 6.44 | AAV.GT5 | $1 \times 10^{12}$ | peripheral vein | neg | neg | 54.98 | |
| #4 | male | K142395 | 10 | 9.1 | AAV.GT5 | $1 \times 10^{12}$ | peripheral vein | neg | 1.58 | 13.41 | hFIX inhibitors detected |
| #5 | male | K132161 | 10 | 7.16 | AAV.GT5 | $1 \times 10^{12}$ | peripheral vein | neg | >10 | 15.9 | |
| #6 | male | K173135 | 9 | 12.75 | AAV-Spark100 | $1 \times 10^{12}$ | peripheral vein | neg | neg | 58.93 | |
| #7 | male | X1003533 | 10 | 10.52 | AAV-Spark100 | $1 \times 10^{12}$ | peripheral vein | neg | neg | 22.93 | hFIX inhibitors detected |
| #8 | male | K173125 | 9 | 9.51 | AAV-Spark100 | $1 \times 10^{12}$ | peripheral vein | neg | neg | 14.38 | |
| #9 | male | X1403793 | 6 | 5.57 | AAV-Spark100 | $1 \times 10^{12}$ | peripheral vein | neg | neg | 117.1 | |
| #59 | male | 1521805011 | 4 | 5.7 | AAV.GT5 | $3 \times 10^{11}$ | intra-hepatic artery | neg | ND | 54.09 | |
| #61 | male | 1521802001 | 4 | 4.52 | AAV.GT5 | $3 \times 10^{11}$ | intra-hepatic artery | neg | ND | 51.27 | |

neg, negative; ND, not determined.

### Supplemental table S2. Microminipigs included in this study

| Number | Sex | Individual number | Date of birth | Body weight (kg) | Vector used | Vector dose (vg/kg) | Administration route | NAb titer against GT5 | NAb titer against Spark100 | MAX of hFIX Ag (%) | Note |
|---|---|---|---|---|---|---|---|---|---|---|---|
| 19-001 | male | 4123 | 2018/11/3 | 12 | AAV-Spark100 | $1 \times 10^{12}$ | portal vein | neg | neg | 8.39 | |
| 19-002 | male | 4126 | 2018/11/4 | 16.6 | AAV-Spark100 | $1 \times 10^{12}$ | portal vein | neg | neg | 2.81 | hFIX inhibitors detected |
| 20-001 | male | 4471 | 2019/12/16 | 15.3 | AAV.GT5 | $1 \times 10^{12}$ | peripheral vein | neg | ND | 15.1 | |
| 20-002 | male | 4480 | 2019/12/19 | 15.7 | AAV.GT5 | $1 \times 10^{12}$ | peripheral vein | neg | ND | 1.2 | |
| 20-003 | male | 4482 | 2020/1/1 | 15.3 | AAV.GT5 | $1 \times 10^{12}$ | portal vein | neg | ND | 101.2 | |
| 20-004 | male | 4501 | 2020/1/9 | 15.8 | AAV.GT5 | $1 \times 10^{12}$ | portal vein | neg | ND | 46.6 | |
| 21-001 | male | 4544 | 2020/4/12 | 17.9 | AAV.GT5 | $1 \times 10^{12}$ | portal vein | neg | ND | 121.3 | |
| 21-002 | male | 4628 | 2021/2/11 | 8.7 | AAV.GT5 | $1 \times 10^{12}$ | peripheral vein | neg | ND | 56.6 | |
| 21-003 | male | 4630 | 2021/2/11 | 6.3 | AAV.GT5 | $1 \times 10^{12}$ | intra-hepatic artery | neg | ND | 86.7 | |
| 21-004 | male | 4632 | 2021/2/12 | 7.7 | AAV.GT5 | $1 \times 10^{12}$ | intra-hepatic artery | neg | ND | 63.4 | |
| 21-005 | male | 4562 | 2020/4/22 | 15.6 | AAV-Spark100 | $1 \times 10^{12}$ | peripheral vein | neg | neg | 122.9 | hFIX inhibitors detected |
| | | | | | AAV.GT5 | | intra-hepatic artery | 15.8 | 394 | 75.2 | NAb against GT5 observed |
| 21-006 | male | 4601 | 2020/11/16 | 8.9 | AAV-Spark100 | $1 \times 10^{12}$ | peripheral vein | neg | neg | 16 | |
| | | | | | AAV.GT5 | | intra-hepatic artery | neg | 7.2 | 369 | |

neg, negative; ND, not determined.

**Supplemental table S3. Oligonucleotide primers for this study**

| | |
|---|---|
| For titration of AAV vectors and qPCR of vg in plasma | |
| SV40_F | 5'-GCAATAGCATCACAAATTTCAC-3' |
| SV40_R | 5'-GATCCAGACATGATAAGATACATTG-3' |
| SV40 Probe | 5'-TCACTGCATTCTAGTTGTGGTTTGTCCA-3' |
| For qPCR of mRNA expression | |
| *F9* CO_F | 5'-GCACAGAGGGCTACAGACTG-3' |
| *F9* CO_R | 5'-CTGGTCAGCTTGCTGGTCTG-3' |
| For qPCR of vg in genomic DNA | |
| *Fluc*_F | 5'-GCGCGGAGGAGTTGTGTT-3' |
| *Fluc*_R | 5'-TCTGATTTTTCTTGCGTCGAGTT-3' |
| *F9* CO_F | 5'-GCACAGAGGGCTACAGACTG-3' |
| *F9* CO_R | 5'-CTGGTCAGCTTGCTGGTCTG-3' |
| For target sequence amplification in next-generation sequencing analysis | |
| NGS *F9*_F | 5'-GCATTGCTGACAAAGAGTACACCA-3' |
| NGS *F9*_R | 5'-ATGATGCCTGTCAGGAAGCT-3' |

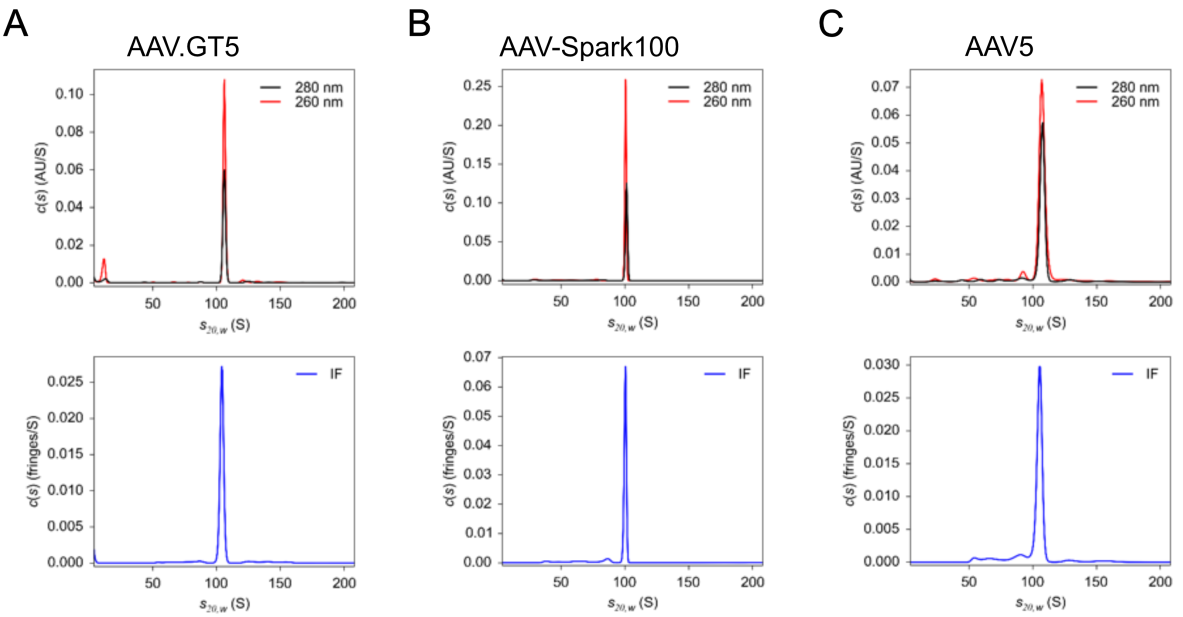

A

| | I | II | Average |
|---|---|---|---|
| EC | 1.2 | 1.7 | 1.45 |
| PP | 3.5 | 5 | 4.25 |
| FP | 89 | 78 | 83.5 |

B

| | I | II | Average |
|---|---|---|---|
| EC | 5.3 | 0 | 2.65 |
| PP | 6.3 | 2.8 | 4.8 |
| FP | 83.7 | 89.1 | 86.4 |

C

| | I | II | Average |
|---|---|---|---|
| EC | 2.2 | 5 | 3.6 |
| PP | 11.7 | 3 | 7.35 |
| FP | 82.1 | 85.6 | 83.85 |

**Supplemental figure S1. The purity of AAV vectors purified by gradient centrifugation with cesium chloride.** The purity of AAV was determined by analytical ultracentrifugation. Representative results of distribution plot of the sedimentation coefficient [UV absorbance (Top) and Rayleigh interference optics (Middle)] of AAV.GT5 (A), AAV-Spark100 (B), and AAV5 (C) harboring human coagulation factor IX with Padua mutation (hFIX-Padua) gene driven by HCRhAAT promoter. Average proportions (%) of the empty capsid (EC), partial particle (PP), and full particle (FP) from two experiments are shown in the table below.

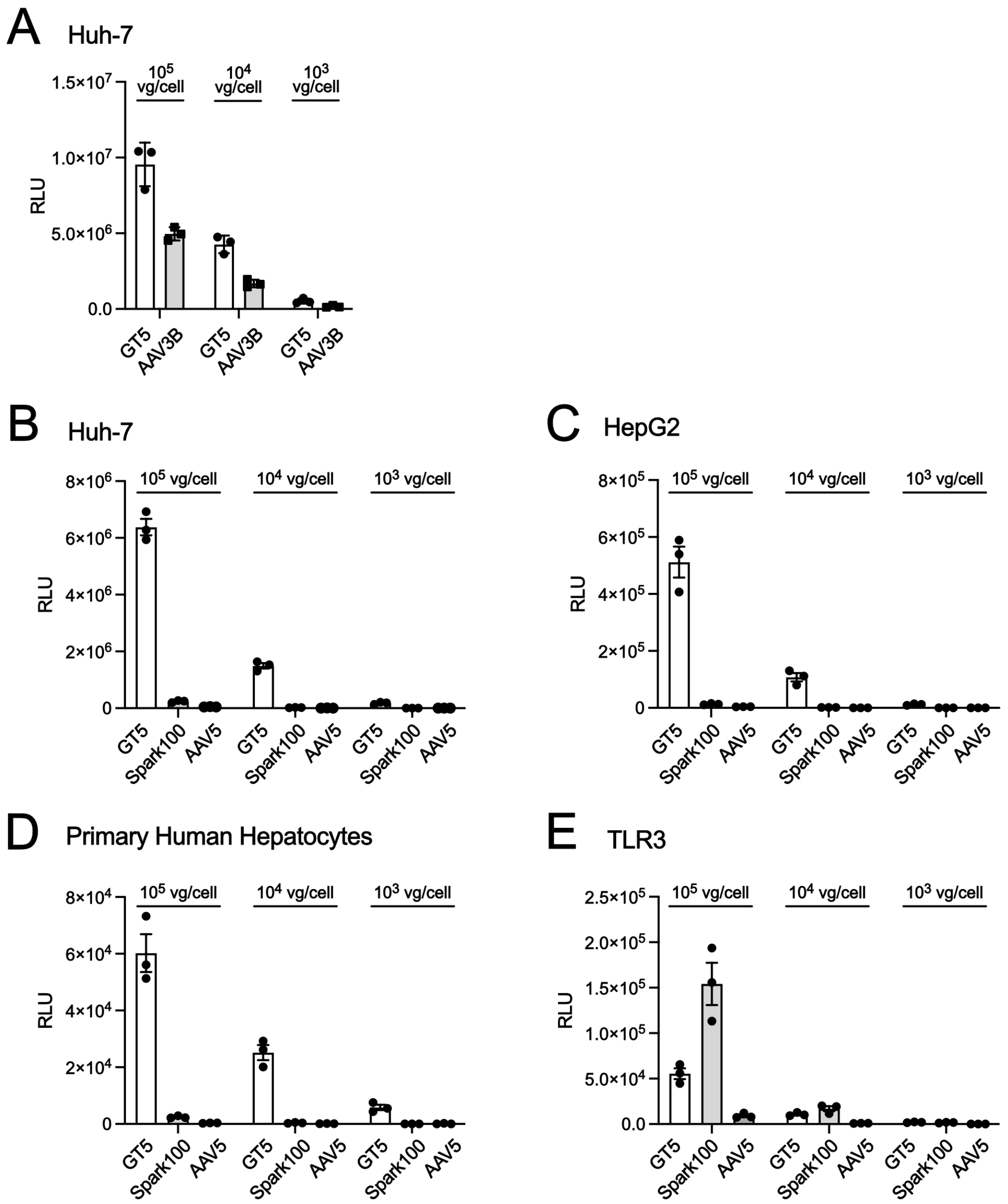

**Supplemental figure S2. Efficient transduction in human hepatocytes by AAV.GT5 *in vitro*.** (A) Huh-7 cells were transduced with AAV.GT5 or AAV3B vector harboring the transgene cassette (CAG promoter, luciferase gene, and SV40 polyA) an indicated multiplicity of infection (MOI). (B–E) AAV.GT5, AAV-Spark100, and AAV5 vectors harboring the same gene cassette transduced Huh-7 cells (B), HepG2 cells (C), primary human hepatocytes (D), and TLR3 cells (E) with an indicated MOI. Luciferase expression in the cell lysates was determined by a luminometer and expressed as a relative light unit (RLU). Luciferase expression in the cell lysates was determined by a luminometer and expressed as RLU. Values are presented as mean ± SEM (n = 3).

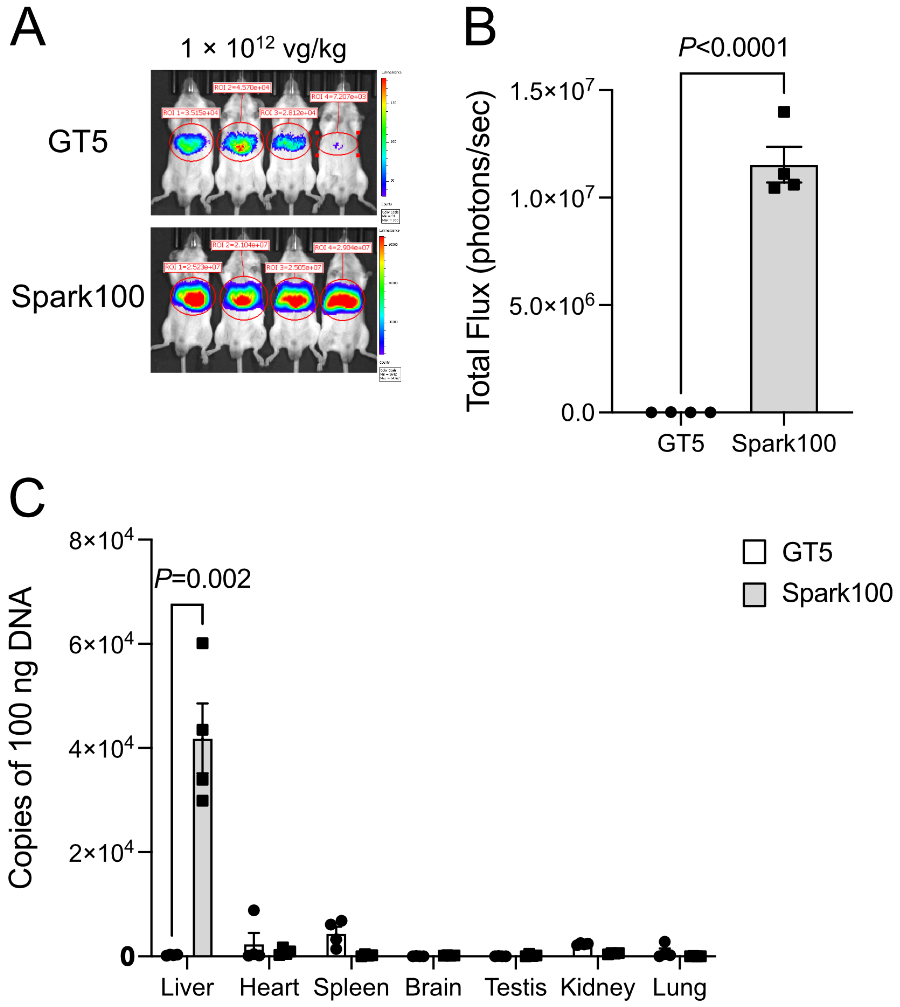

**Supplemental figure S3. The transgene expression in the liver in SCID mice by intravenous injection of AAV.GT5 and AAV-Spark100.** AAV.GT5 or AAV-Spark100 harboring luciferase gene driven by CAG promoter were intravenously injected into SCID mice (1 × $10^{12}$ vg/kg). (A, B) Photons transmitted through the body were analyzed using an IVIS Imaging System. (A) Representative data of IVIS imaging system at 4 weeks after the vector administration. (B) Quantitative data of each mouse were expressed as photon units (photons/second) (n = 4). Values are presented as mean ± SEM (n = 4). (C) AAV genome in organs was determined by quantitative PCR at 4 weeks after the vector administration (n = 4 in each group). Transduced values are mean ± SEM (n = 4). Statistical analysis between the two groups was analyzed by Student *t*-test. Actual *P* values are described in the figure.

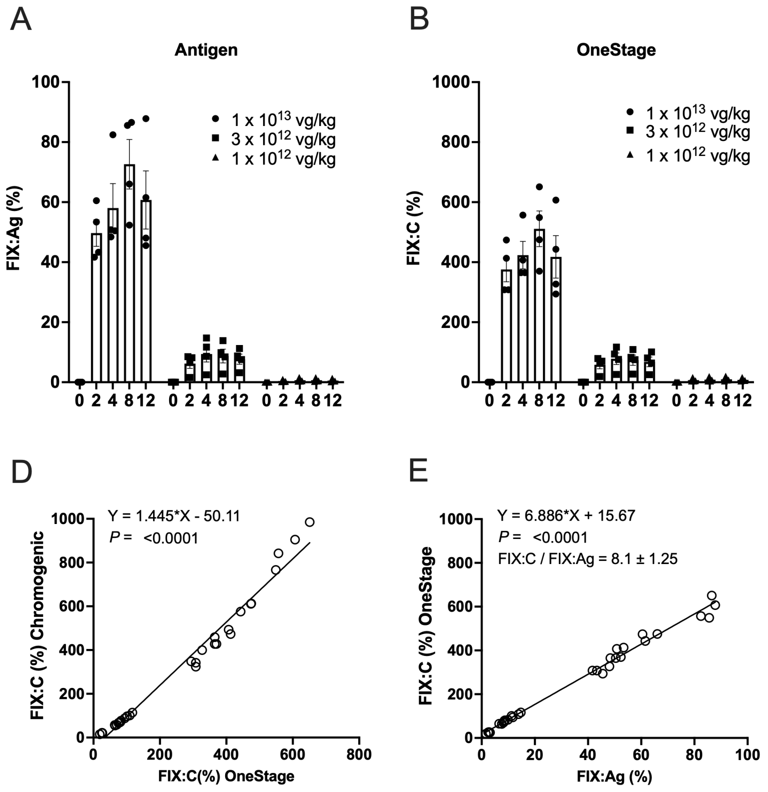

**Supplemental figure S4. The increase in plasma FIX in FIX-deficient mice by intravenous injection of AAV.GT5.** AAV.GT5 harboring human coagulation factor IX with Padua mutation (hFIX-Padua) gene driven by HCRhAAT promoter were intravenously injected into Factor IX-deficient mice ($1 \times 10^{12}$, $3 \times 10^{12}$, or $1 \times 10^{13}$ vg/kg). The increase in FIX antigen (FIX:Ag) (A) and FIX activity (FIX:C) measured by one-stage clotting assay (B) and chromogenic assay (C) are shown. Values are presented as mean ± SEM (n = 4). (D, E, and F) The correlation between the two measurements in plasma is shown. (D) FIX:C (one-stage clotting assay) and FIX:C (chromogenic assay). (E) FIX:Ag and FIX:C (one-stage clotting assay). (F) FIX:Ag and FIX:C (chromogenic assay).

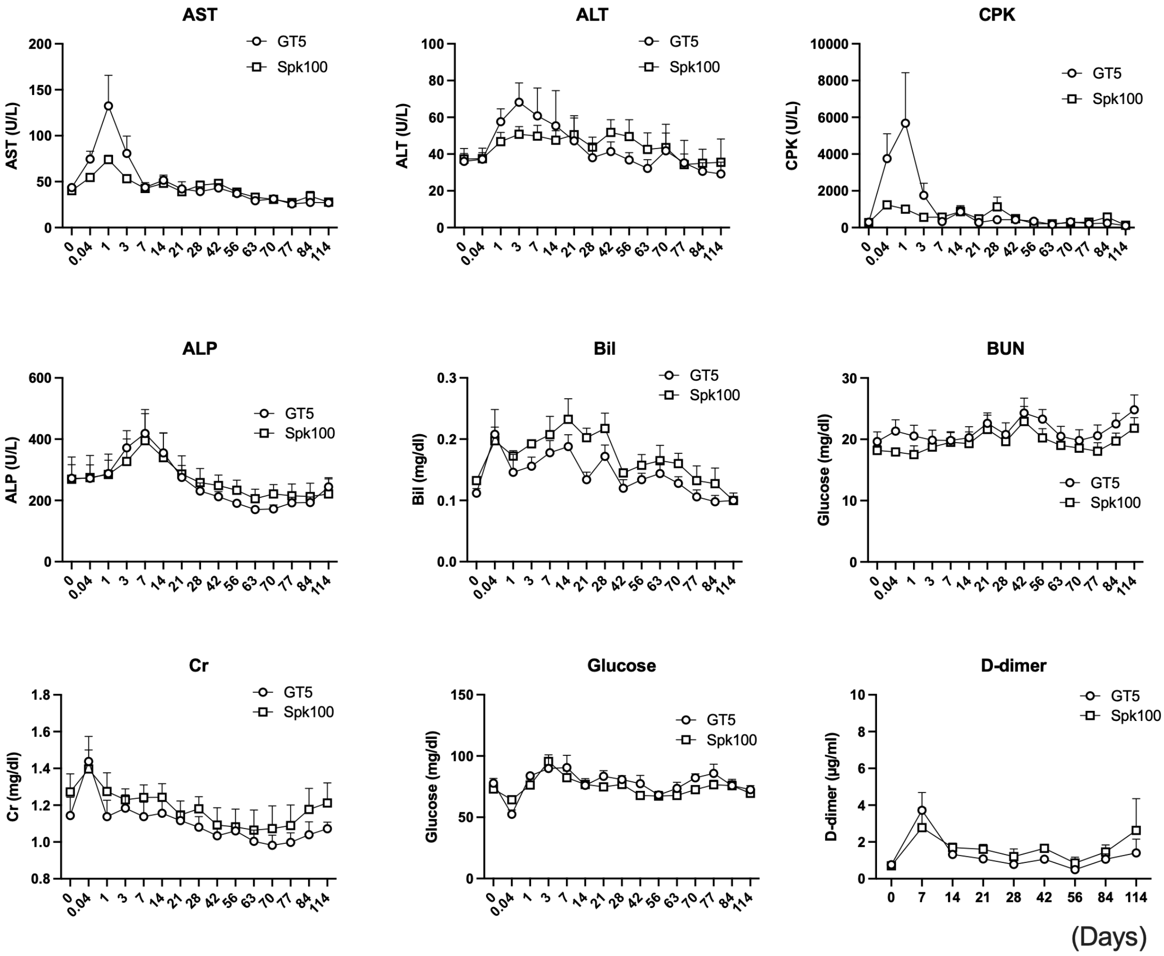

**Supplemental figure S5. Changes in laboratory test parameters after AAV vector administration in cynomolgus monkeys.** AAV.GT5 or AAV-Spark100 harboring human coagulation factor IX with Padua mutation (hFIX-Padua) gene driven by HCRhAAT promoter were intravenously injected into cynomolgus monkeys ($1 \times 10^{12}$ vg/kg). The changes in serum aspartate aminotransferase (AST), alanine aminotransferase (ALT), creatine kinase (CPK), alkaline phosphatase (ALP), total bilirubin (Bil), blood urea nitrogen (BUN), creatinine (Cr), blood glucose (Glucose) and plasma D-dimer were measured. Values are presented as mean ± SEM (n = 4).

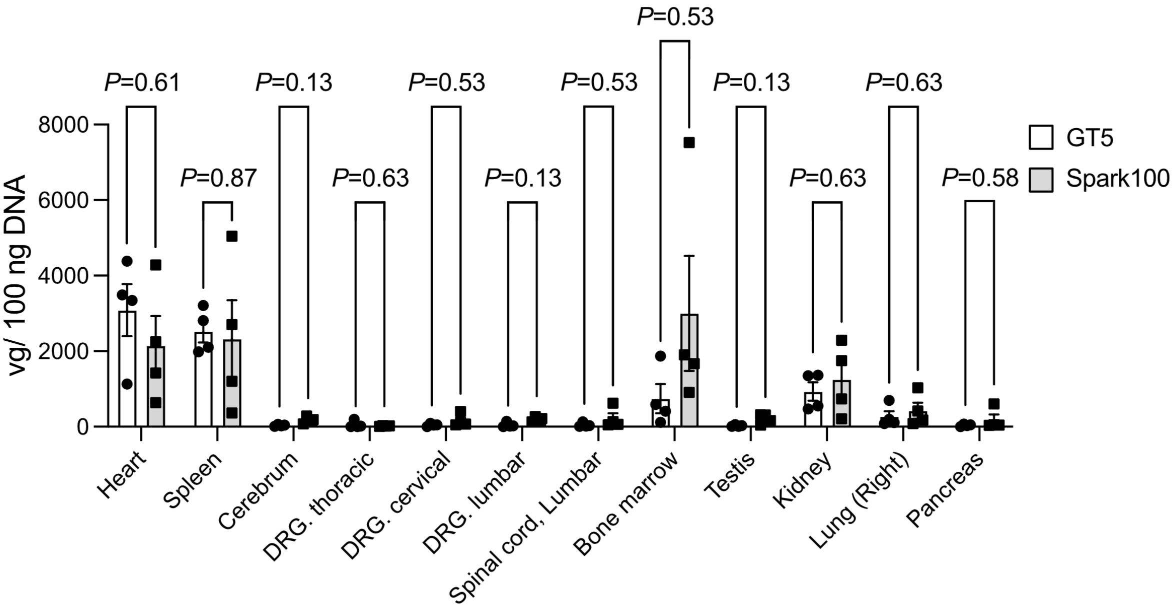

**Supplemental figure S6. AAV genome in cynomolgus monkeys by intravenous injection of AAV.GT5 and AAV-Spark100.** AAV.GT5 or AAV-Spark100 harboring human coagulation factor IX with Padua mutation (hFIX-Padua) gene driven by HCRhAAT promoter were intravenously injected into cynomolgus monkeys ($1 \times 10^{12}$ vg/kg). AAV genome in organs without a liver was determined by quantitative PCR at 114 days after the vector administration. Values are presented as mean ± SEM (n = 4). Statistical analysis between the two groups was performed by Student *t*-test. Actual *P* values are described in the figure.
DRG, dorsal root ganglion.

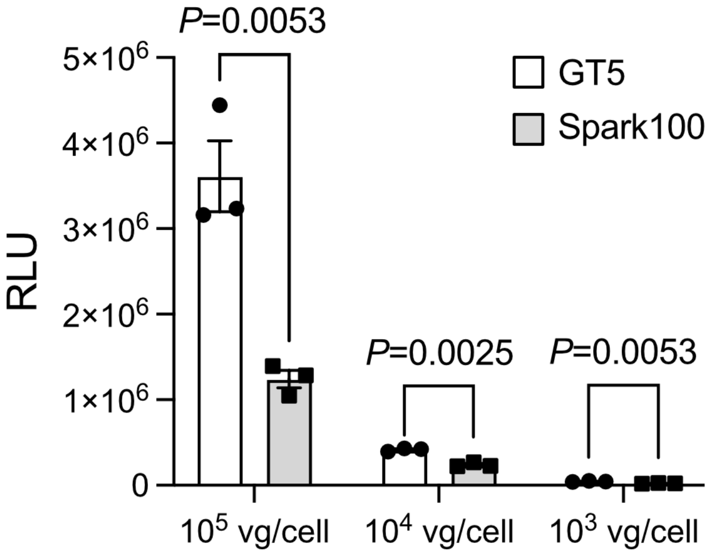

**Supplemental figure S7. Transduction efficacy with AAV.GT5 and AAV-Spark100 in porcine hepatocyte.** AAV.GT5 and AAV-Spark100 vectors harboring the transgene gene cassette (CAG promoter, luciferase gene, and SV40 polyA) were transduced primary porcine hepatocytes at an indicated multiplicity of infection (MOI). Luciferase expression in the cell lysates was determined by a luminometer and expressed as a relative light unit (RLU). Values are expressed as mean ± SEM (n = 3).

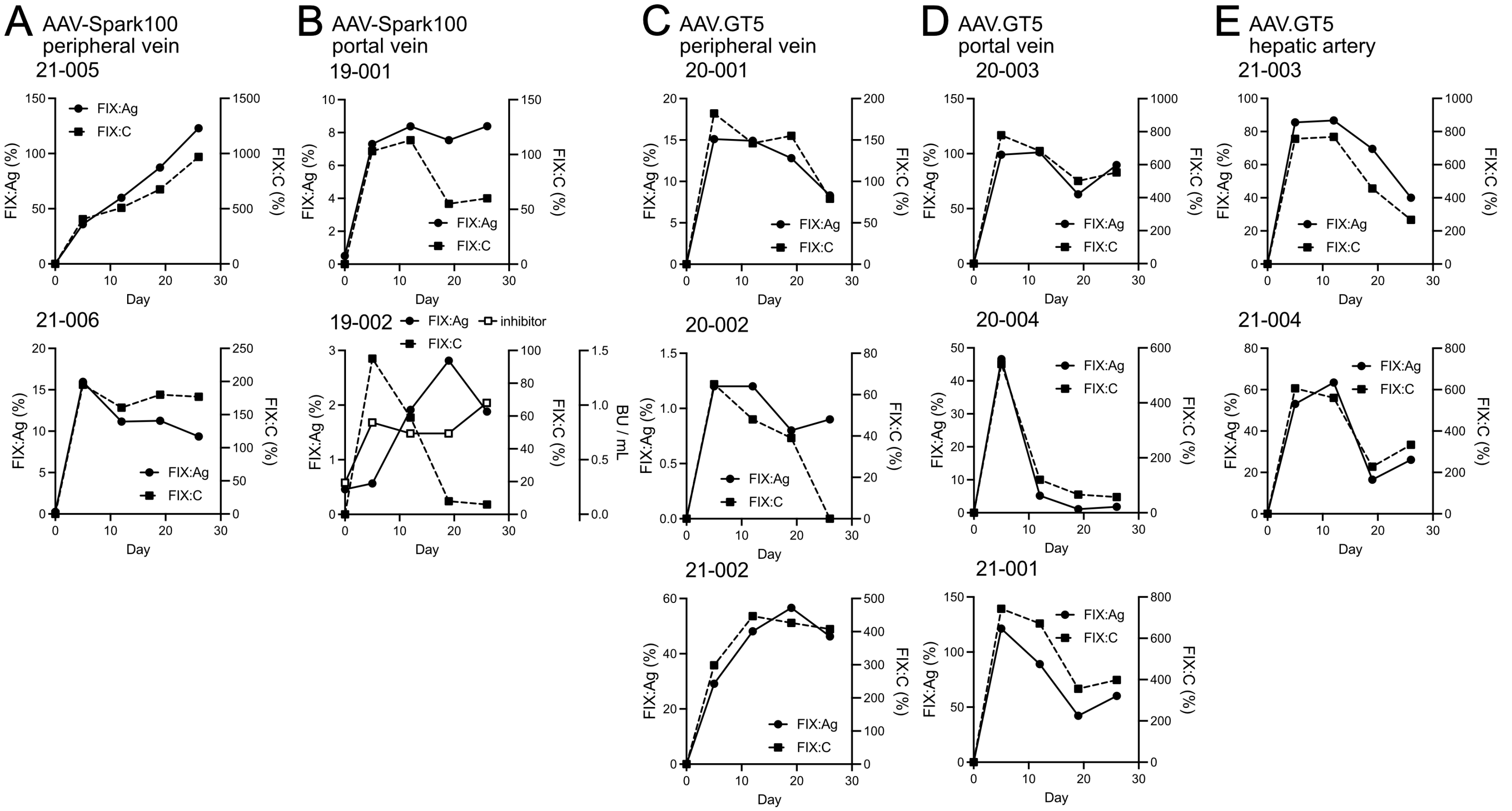

**Supplemental figure S8. Transduction efficacy with AAV.GT5 and AAV-Spark100 in pigs.** AAV.GT5 (A, B) or AAV-Spark100 harboring human coagulation factor IX with Padua mutation (hFIX-Padua) gene driven by HCRhAAT promoter were administrated into male microminipigs through a peripheral vein, left portal vein, or hepatic artery ($1 \times 10^{12}$ vg/kg). (A–E) The changes in plasma FIX:Ag and FIX:C after the administration in each pig are shown. FIX neutralizing antibody was developed in one pig (#19-002 in AAV-Spark100).

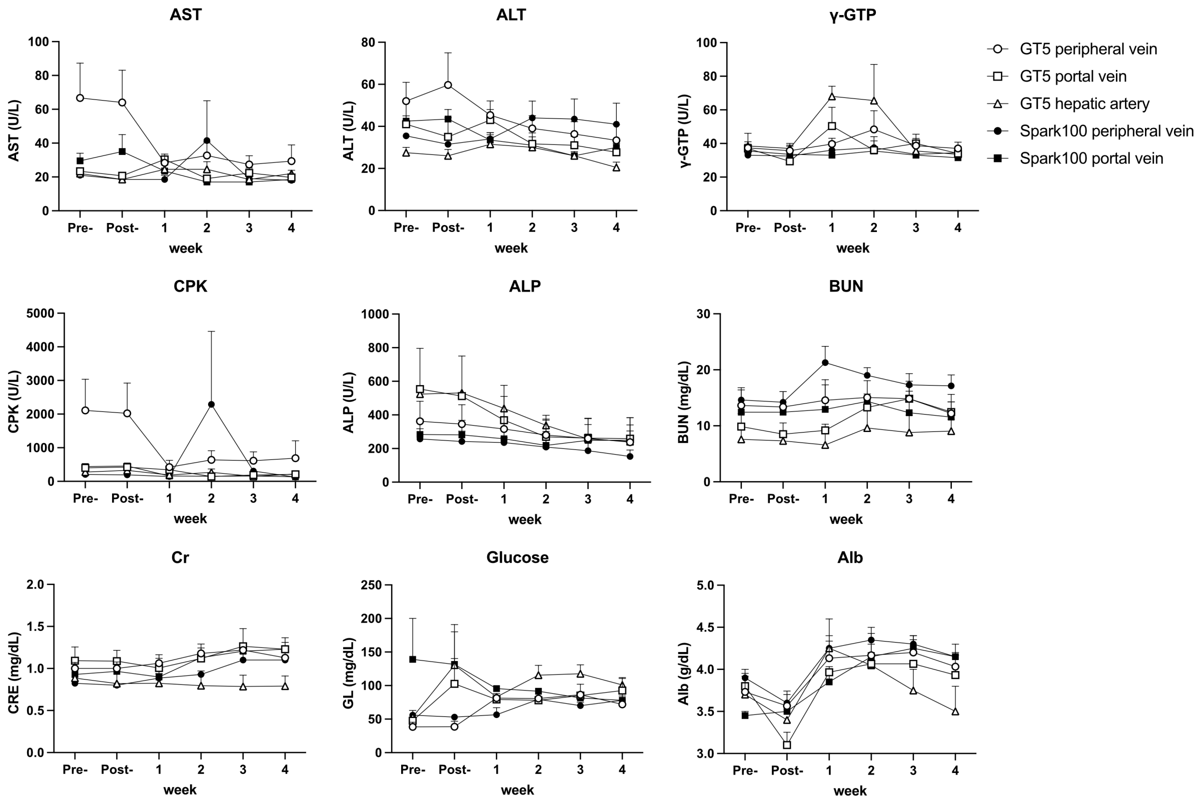

**Supplemental figure S9. Changes in laboratory test parameters after AAV vector administration in pigs.** AAV.GT5 or AAV-Spark100 harboring human coagulation factor IX with Padua mutation (hFIX-Padua) gene driven by HCRhAAT promoter were injected into male microminipigs ($1 \times 10^{12}$ vg/kg). The changes in serum aspartate aminotransferase (AST), alanine aminotransferase (ALT), γ-glutamyl transpeptidase (γ-GTP), creatine kinase (CPK), alkaline phosphatase (ALP), blood urea nitrogen (BUN), creatinine (Cr), blood glucose (Glucose) and Albumin (Alb) were measured. Values are presented as mean ± SEM (AAV-Spark100: n = 2, AAV.GT5: n = 3 in peripheral vein and left portal vein; n = 2 in hepatic artery).

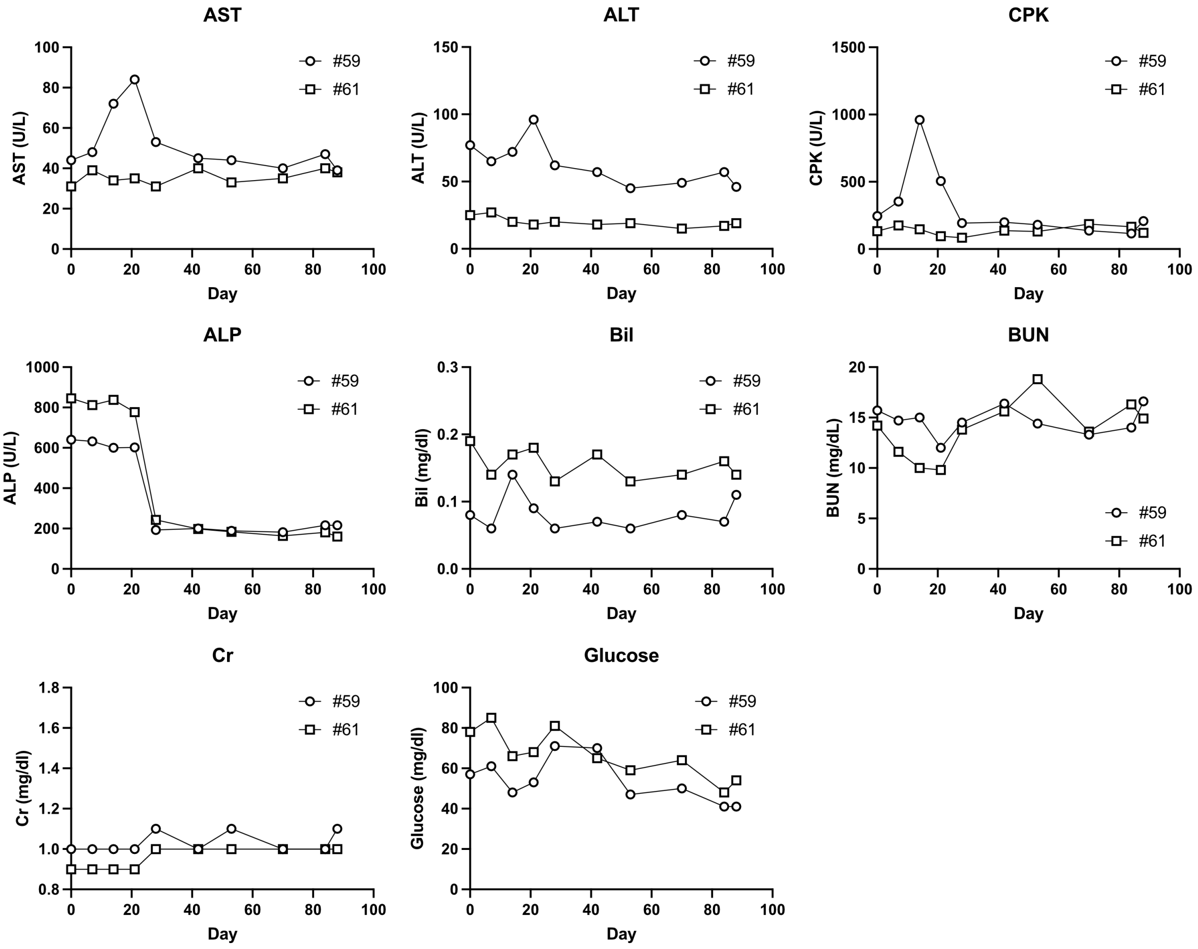

**Supplemental figure S10. Changes in laboratory test parameters after AAV vector administration via a hepatic artery in cynomolgus monkeys.** AAV.GT5 harboring human coagulation factor IX with Padua mutation (hFIX-Padua) gene driven by HCRhAAT promoter were injected into cynomolgus monkey through the hepatic artery ($3 \times 10^{11}$ vg/kg). The changes in serum aspartate aminotransferase (AST), alanine aminotransferase (ALT), creatine kinase (CPK), alkaline phosphatase (ALP), total bilirubin (Bil), blood urea nitrogen (BUN), creatinine (Cr), and blood glucose (Glucose) were measured. The data of each monkey is shown.

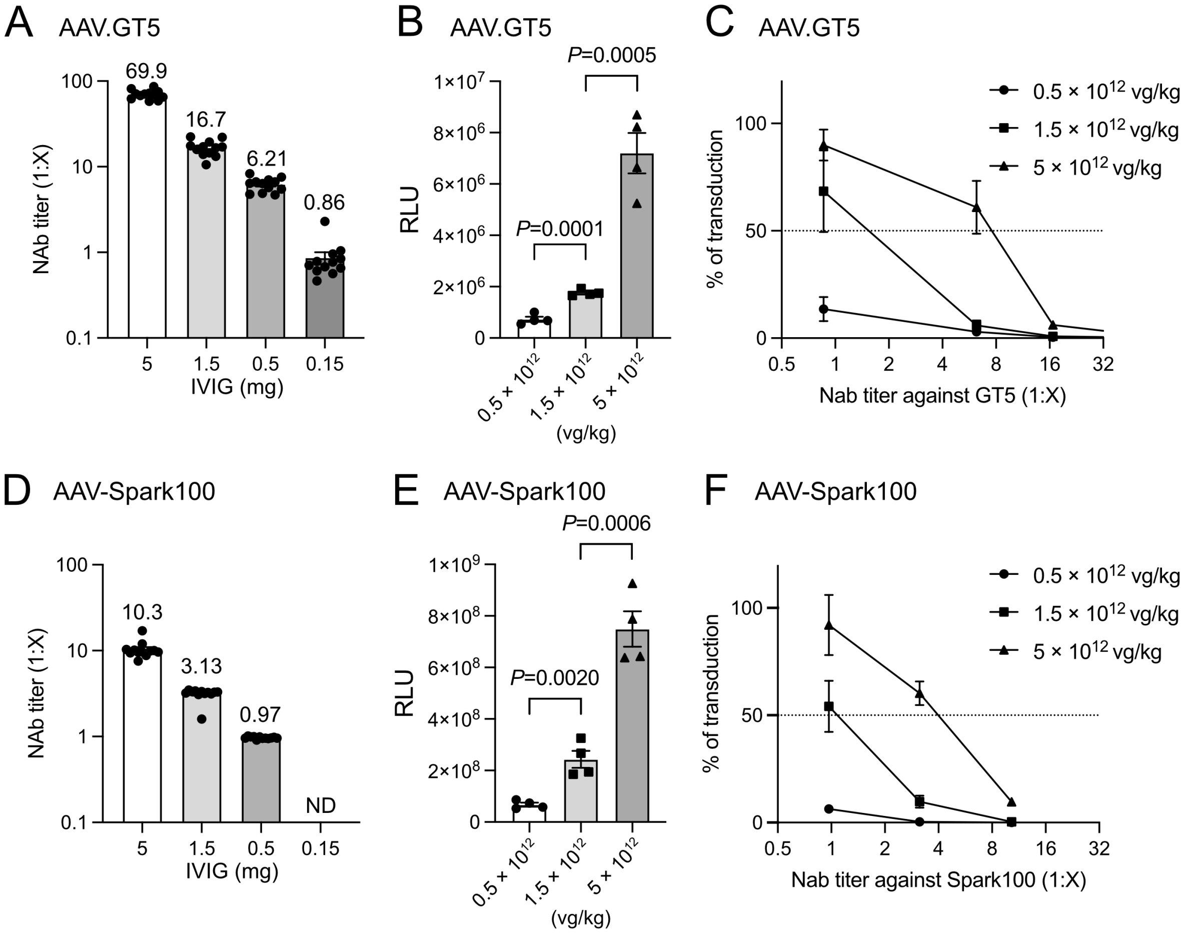

**Supplemental figure S11. The threshold of neutralizing antibody (NAb) to inhibit *in vivo* transduction of AAV.GT5 and AAV-Spark100.** An indicated dose of human immunoglobulin (IVIG) was intravenously administered to C57BL6/J mice. Phosphate-buffered saline was used in a control experiment. Blood was drawn 1 hour after the IVIG injection and then the AAV.GT5 (B and C) or AAV-Spark100 (E and F) vector expressing secNanoLuc ($0.5 \times 10^{12}$ vg/kg; $1.5 \times 10^{12}$ vg/kg; $5 \times 10^{12}$ vg/kg) was administrated to the mice. (A, D) Serum NAb titers against AAV.GT5 (A) or AAV-Spark100 (D) after IVIG injection. Values are presented as mean ± SEM (n = 12). (B, E) Serum secNanoLuc activity in mice treated with an indicated dose of AAV.GT5 (B) or AAV-Spark1000 (E) 30 days after the vector injection. Statistical analysis was analyzed by Student *t*-test. Actual *P* values are described in the figure. (C, F) Inhibition of *in vivo* liver transduction with an indicated dose of AAV.GT5 (C) or AAV-Spark1000 (F) by the pretreatment with IVIG. The horizontal axis indicates NAb titer after IVIG administration at a dose described in A and D. Values are presented as mean ± SEM (n = 4).

ND, not determined.

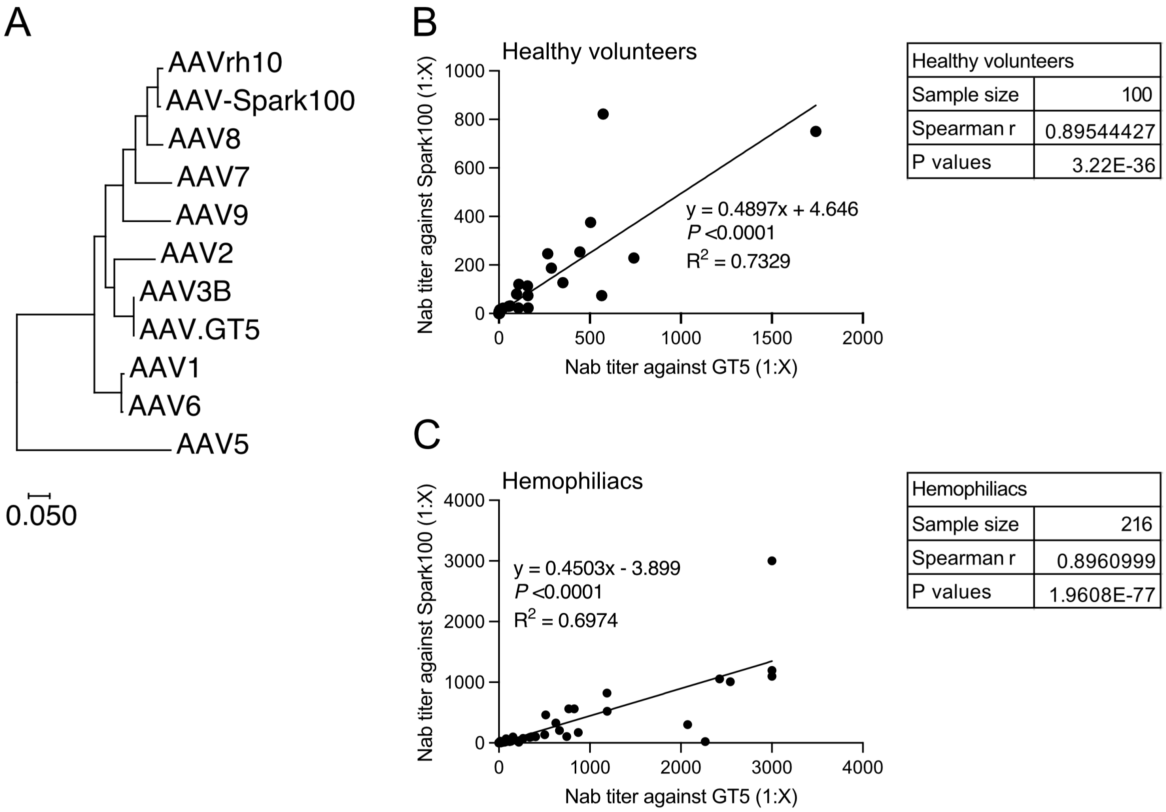

**Supplemental figure S12. Correlation between NAb titers against AAV.GT5 and AAV-Spark100.** (A) The topology is based on the similarity of the capsid sequences of various AAV serotypes, including AAV.GT5 and AAV-Spark100. (B, C) The correlation between NAb titers against AAV.GT5 and AAV-Spark100 in healthy volunteers (B) and hemophiliacs (C). Statistical analysis between the two groups was analyzed by Spearman's rank correlation coefficient. Spearman r values and *P* values are described in the figure.

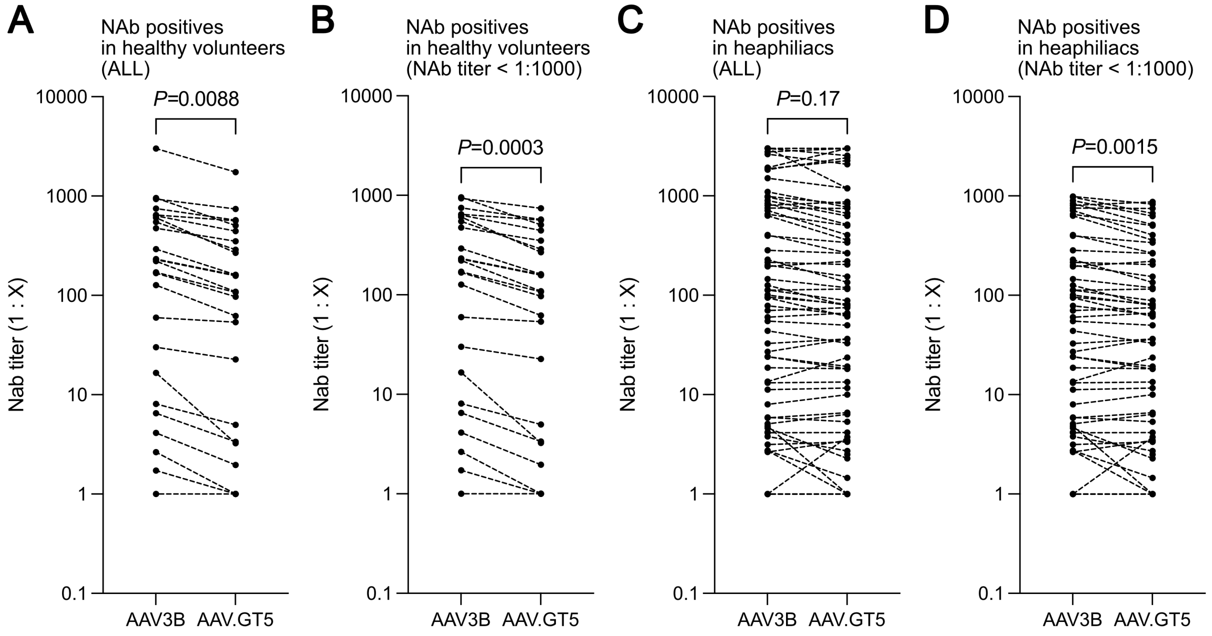

**Supplemental figure S13. Comparison of NAb titers between AAV3B and AAV.GT5 vector in the Japanese population.** (A, B) The NAb titer against AAV3B and AAV.GT5 in all healthy volunteers (A) and healthy volunteers with NAb titers less than 1:1000 (B) (C, D) The NAb titer against AAV3B and AAV.GT5 in all hemophiliacs (C) and hemophiliacs with NAb titers less than 1:1000 (D). Statistical analysis was performed by Paired Student *t*-test. Actual *P* values are described in the figure. Data on NAb titers against AAV3B in the Japanese population were taken from our previously reported study.[2]

**Video 1 Representative contrast images before the intra-portal administration in pig.** Portal vein (left image), left branch of the portal vein (Right image).

**Video 2 Representative contrast images before th**e intra-hepatic artery **administration in pig.** Common hepatic artery (left image), proper hepatic artery (Right image).

**Video 3 Representative contrast images before the intra-hepatic artery administration in macaque.**
